# *Ex-Vivo* Equine Cartilage Explant Osteoarthritis Model - A Metabolomics and Proteomics Study

**DOI:** 10.1101/2020.03.03.974501

**Authors:** James R Anderson, Marie M Phelan, Laura Foddy, Peter D Clegg, Mandy J Peffers

**Author notes:** Corresponding author email address James R Anderson.

## Abstract

Osteoarthritis (OA) is an age-related degenerative musculoskeletal disease characterised by loss of articular cartilage, synovitis, abnormal bone proliferation and subchondral bone sclerosis. Underlying OA pathogenesis is yet to be fully elucidated with no OA specific biomarkers in clinical use. *Ex-vivo* equine cartilage explants (n=5) were incubated in TNF-α/IL-1β supplemented culture media for 8 days, with media removed and replaced at 2, 5 and 8 days. Acetonitrile metabolite extractions of 8 day cartilage explants and media samples at all time points underwent 1D ^1^H nuclear magnetic resonance metabolomic analysis with media samples also undergoing mass spectrometry proteomic analysis. Within the cartilage, metabolites glucose and lysine were elevated following TNF-α/IL-1β treatment whilst adenosine, alanine, betaine, creatine, myo-inositol and uridine levels decreased. Within the culture media, four, four and six differentially abundant metabolites and 154, 138 and 72 differentially abundant proteins, with > 2 fold change, were identified for 1-2 day, 3-5 day and 6-8 day time points respectively. Nine potential novel OA neopeptides were elevated in treated media. Our innovative study has identified differentially abundant metabolites, proteins and extracellular matrix derived neopeptides, providing insightful information on OA pathogenesis, enabling potential translation for clinical markers and possible new therapeutic targets.

## Introduction

Osteoarthritis (OA) is an age-related degenerative musculoskeletal disease characterised by loss of articular cartilage, synovial membrane dysfunction, abnormal bone proliferation, subchondral bone sclerosis and altered biochemical and biomechanical properties ^1, 2^. For horses in the UK, OA is one of the leading welfare issues, resulting in substantial morbidity and mortality ^3, 4^. It is estimated that OA accounts for 60% of lameness seen in horses ^5^. Within OA, extracellular matrix (ECM) degradation is driven by multiple matrix metalloproteinases (MMPs) and a disintegrin and metalloproteinases with thrombospondin motifs (ADAMTSs) ^6^. However, the underlying pathogenesis of OA is yet to be fully elucidated with no disease-modifying treatments currently available ^7, 8^. Whilst a number of putative biomarkers have been identified for OA diagnosis in the horse, none are currently used within clinical practice ^9^. Presently, equine OA is predominantly diagnosed through diagnostic imaging and clinical examination. However, due to the slow onset of the condition, this often leads to substantial pathology of the joint, particularly to articular cartilage prior to diagnosis ^10^. There is therefore a need to develop diagnostic tests which are sensitive and specific to the early stages of OA, which are repeatable and reproducible, as well as gaining a greater understanding of the underlying pathogenesis ^11, 12^. Early detection of OA could enable timely management interventions which could potentially slow the progression of the disease.

Tumour necrosis factor-α (TNF-α) and interleukin-1β (IL-1β) are both pro-inflammatory cytokines which are central in OA pathogenesis ^13^. TNF-α and IL-1β are secreted by mononuclear cells, synoviocytes and articular cartilage and upregulate gene expression of MMPs, ADAMTS-4 and ADAMTS-5, leading to significant ECM degradation ^14–16^. Elevations in TNF-α and IL-1β are regularly identified within OA synovial fluid, including that of horses ^17–19^. TNF-α and IL-1β have therefore become established experimental treatments for modelling OA pathology within *in vitro* and *ex-vivo* studies, having been used both independently and as a combined treatment ^20–26^.

Proteomics is the systematic, large scale study of proteins within biological systems to assess quantities, isoforms, modifications, structure and function ^27^. Previous studies have undertaken mass spectrometry (MS) based proteomics using TNF-α and IL-1β OA models for secretome analysis of chondrocytes *in vitro* and *ex-vivo* cartilage explants ^20–22, 25^. Results from these studies included increased media levels of MMPs, cartilage oligomeric matrix protein (COMP), aggrecan and collagen VI.

During OA pathology, disease-associated peptide fragments (neopeptides) are generated from cartilage breakdown due to increased enzymatic activity/abundance of MMPs, ADAMTSs, cathepsins and serine proteases ^28–30^. MS analysis of these neopeptides can then be applied to identify potential early OA biomarkers ^31^. Previously a murine 32 amino acid peptide fragment, generated through increased activity of MMP and ADAMTS-4/5 and subsequent aggrecan degradation, was found to drive OA pain via Toll-like receptor 2 ^32^. Neopeptide targeting therefore has the potential to provide a localised analgesic at the site of joint degeneration ^31^. Numerous equine OA studies investigating both synovial fluid (SF) and cartilage have identified potential neopeptides of interest ^28, 33–35^. Development of antibodies targeted to OA specific neopeptides would provide the ability to monitor cartilage degeneration, assess therapeutic response and potentially provide future novel therapeutic targets ^31, 36^.

Metabolomics uses a systematic methodology to comprehensively identify and quantify the metabolic profiles of biological samples ^37^. ^1^H Nuclear magnetic resonance (NMR) metabolomics analysis provides a high level of technical reproducibility with a minimal level of sample preparation ^38^. ^1^H NMR analysis has previously been used to investigate OA in the SF of humans, horses, pigs and dogs ^9, 39–45^. Synovial metabolites alanine, choline, creatine and glucose have been identified as differentially abundant in OA across multiple studies and species ^9, 39, 41–44^. NMR techniques have also previously been used to characterise cartilage with high resolution magical angle spinning (HRMAS) NMR utilised to assess enzymatic degradation of bovine cartilage ^46–48^. A guinea pig OA model using HRMAS NMR identified elevations in methylene resonances associated with chondrocyte membrane lipids and an increase in mobile methyl groups of collagen ^49^. Another HRMAS NMR study of human OA cartilage identified a reduction in alanine, choline, glycine, lactate methyne and N-acetyl compared to healthy control cartilage ^50^. However, no NMR studies to date have investigated the metabolic profile of culture media following the incubation of *ex-vivo* cartilage within an OA model.

This is the first study to carry out ^1^H NMR metabolomic analysis of extracted cartilage metabolites and also to undertake ^1^H NMR analysis of culture media using the TNF-α/IL-1β *ex-vivo* OA cartilage model. Additionally, this is also the first study to use a multi ‘omics’ approach to simultaneously investigate the metabolomic profile of *ex-vivo* cartilage and metabolomic/proteomic profiles of culture media using this OA model. It was hypothesised that following TNF-α/IL-1β treatment of *ex-vivo* equine cartilage, ^1^H NMR metabolomic and MS proteomic platforms would identify a panel of cartilage metabolites which were able to differentiate control from treated cartilage and a panel of metabolites, proteins and neopeptides within the associated culture media which were differentially abundant at each tested time point of the OA model.

## Methods

### Equine *Ex-Vivo* Cartilage Collection

Full thickness cartilage was removed from all articular surfaces within five separate metacarpophalangeal joints of five nine-year-old mares of unknown breed within 24 hr of slaughter at a commercial abattoir (F Drury and Sons, Swindon, UK). Cartilage samples were collected as a by-product of the agricultural industry. The Animals (Scientific Procedures) Act 1986, Schedule 2, does not define collection from these sources as scientific procedures and ethical approval was therefore not required. Cartilage collected from all joints was considered macroscopically normal with a score of 0 according to the OARSI histopathology initiative scoring system for horses ^51^ (Figure S1). Cartilage was washed in complete media containing Dulbecco’s modified Eagle’s medium (DMEM, 31885-023, Life Technologies, Paisley, UK) supplemented with 10% (v/v) foetal calf serum (FCS, Life Technologies), 5 μg/ml Amphotericin B (Life Technologies), 100 U/ml Streptomycin and 100 U/ml Penicillin (Sigma-Aldrich, Gillingham, UK) (Figure S2).

Cartilage was dissected into 3 mm^2^ sections and divided into two for each donor (control and treatment wells) on a twelve well plate (Greiner Bio-One Ltd., Stonehouse, UK). Explants were incubated for 24 hr in complete media within a humidified atmosphere of 5% (v/v) CO_2_ at 37°C. Culture media was removed, explants washed in phosphate buffered saline (PBS, Sigma-Aldrich) and replaced with serum free media (control) or serum free media supplemented with 10 ng/ml TNF-α (PeproTech EC Ltd., London, UK) and 10 ng/ml IL-1β (R&D Systems Inc., Minneapolis, Minnesota, USA) (treatment). After 48 hr, media was removed, centrifuged at 13,000*g*, 4°C for 10 min, supernatant removed and ethylenediaminetetraacetic acid (EDTA)-free protease inhibitor cocktail (Roche, Lewes, UK) added to cell-free media. Supernatant was then snap frozen in liquid nitrogen and stored at −80°C. Cartilage was washed in PBS and control/treatment culture media replaced as appropriate. Media collection was repeated at five and eight days. After day eight, cartilage was washed in PBS, weighed, snap frozen in liquid nitrogen and stored at −80°C.

## NMR Metabolomics

### Cartilage - Metabolite Extraction

Equal masses of cultured cartilage explants (in addition to three macroscopically normal equine cartilage samples, each divided into three to assess metabolite extraction reproducibility) were thawed out over ice and added to 500 µl of 50:50 (v/v) ice cold acetonitrile (ThermoFisher Scientific, Massachusetts, USA):dd ^1^H_2_O and incubated on ice for 10 min. Samples were then sonicated using a microtip sonicator at 50 KHz and 10 nm amplitude in an ice-bath for three 30 s periods, interspersed with 30 s rests (that ensured extraction mixture temperature did not exceed 15°C). The extraction mixture was then vortexed for 1 min, centrifuged at 12,000*g* for 10 min at 4°C, supernatant transferred before being snap frozen in liquid nitrogen, lyophilised and stored at −80°C ^37^.

### Cartilage - NMR Sample Preparation

Each lyophilised sample was dissolved through the addition of 200 µl of 100 µM PO_4_^3-^ pH 7.4 buffer (Na_2_HPO_4_, VWR International Ltd., Radnor, Pennsylvania, USA and NaH_2_PO_4_, Sigma-Aldrich), containing 100 µM d4 trimethylsilyl propionate (TSP, Sigma-Aldrich) and 1.2 µM sodium azide (NaN_3_, Sigma-Aldrich) in 99.9% deuterium oxide (^2^H_2_O, Sigma-Aldrich). Samples were vortexed for 1 min, centrifuged at 12,000*g* for 2 min, 190 µl of supernatant removed and transferred into 3 mm outer diameter NMR tubes using a glass pipette.

### Culture Media - NMR Sample Preparation

Culture media was thawed over ice and centrifuged for 5 min at 21,000*g* and 4°C. 150 µl of thawed culture media was diluted to a final volume containing 50% (v/v) culture media, 40% (v/v) dd ^1^H_2_O, 10% ^2^H_2_O and 0.0025% (v/v) NaN_3_, within an overall concentration of 500 mM PO_4_^3-^ pH 7.4 buffer. Samples were vortexed for 1 min, centrifuged at 13,000*g* for 2 min at 4°C, 250 µl supernatant removed and transferred to 3 mm outer diameter NMR tubes using a glass pipette.

### NMR Acquisition

For each individual sample, 1D ^1^H NMR spectra, with the application of a Carr-Purcell-Meiboom-Gill (CPMG) filter to attenuate macromolecule (e.g. proteins) signals, were acquired using the standard vendor pulse sequence cpmgpr1d on a 700 MHz NMR Bruker Avance III HD spectrometer with associated TCI cryoprobe and chilled Sample-Jet autosampler. All spectra were acquired at 25°C, with a 4 s interscan delay, 256 transients for cartilage spectra and 128 transients for media spectra, with a spectral width of 15 ppm. Topsin 3.1 and IconNMR 4.6.7 software programmes were used for acquisition and processing undertaking automated phasing, baseline correction and a standard vendor processing routine (exponential window function with 0.3 Hz line broadening). In addition to all cartilage extract and culture media samples, protease inhibitor cocktail and treatment cytokines TNF-α and IL-1β were also analysed separately to evaluate their metabolite profiles.

### Metabolite Annotation and Identification

All acquired spectra were assessed to determine whether they met minimum reporting standards (as outlined by the Metabolomics Society) prior to inclusion for statistical analysis ^52^. These included appropriate water suppression, flat spectral baseline and consistent linewidths. Metabolite annotations and relative abundances were carried out using Chenomx NMR Suite 8.2 (330-mammalian metabolite library). When possible, metabolite identifications were confirmed using 1D ^1^H NMR in-house spectral libraries of metabolite standards. All raw 1D ^1^H NMR spectra, together with annotated metabolite HMDB IDs and annotation level, are available within the EMBL-EBI MetaboLights repository (www.ebi.ac.uk/metabolights/MTBLS1495) ^53^. Quantile plots of 1D ^1^H NMR spectra are shown in Figure S3.

## Culture Media Proteomics

### Protein Assay and StrataClean^TM^ Resin Processing

Culture media was thawed over ice and centrifuged for 5 min at 21,000*g* and 4°C. Media sample concentrations were determined using a Pierce® 660 nm protein assay (Thermo Scientific, Waltham, Massachusetts, USA). 50 µg of protein for each sample was diluted with dd H_2_O, producing a final volume of 1 ml. StrataClean^TM^ resin (10 µl) (Agilent, Santa Clara, California, USA) was added to each sample, rotated for 15 min, centrifuged at 400*g* for 1 min and the supernatant removed and discarded. Samples were then washed through the addition of 1 ml of ddH_2_O, vortexed for 1 min, centrifuged at 400*g* for 1 min and the supernatant removed and discarded. The wash step was repeated two further times.

### Protein Digestion

160 µl of 25 mM ammonium bicarbonate (Fluka Chemicals Ltd., Gillingham, UK) containing 0.05% (w/v) RapiGest (Waters, Elstree, Hertfordshire, UK) was added to each sample and heated at 80°C for 10 min. DL-Dithiothreitol (Sigma-Aldrich) was added to produce a final concentration of 3 mM, incubated at 60°C for 10 min then iodoacetamide (Sigma-Aldrich) added (9 mM final concentration) and incubated at room temperature in the dark for 30 min. 2 µg of proteomics grade trypsin (Sigma-Aldrich) was added to each sample, rotated at 37°C for 16 hr and trypsin treatment then repeated for a 2 hr incubation. Samples were centrifuged at 1,000*g* for 1 min, digest removed, trifluoroacetic acid (TFA, Sigma-Aldrich) added (0.5% (v/v) final concentration) and rotated at 37°C for 30 min. Finally, digests were centrifuged at 13,000*g* and 4°C for 15 min and the supernatant removed and stored at 4°C.

### Label Free LC-MS/MS

All media digests were randomised and individually analysed using LC-MS/MS on an UltiMate 3000 Nano LC System (Dionex/Thermo Scientific) coupled to a Q Exactive^TM^ Quadrupole-Orbitrap instrument (Thermo Scientific). Full LC-MS/MS instrument methods are described in the supporting information. Tryptic peptides, equivalent to 250 ng of protein, were loaded onto the column and run over a 1 hr gradient, interspersed with 30 min blanks (97% (v/v) high performance liquid chromatography grade H_2_0 (VWR International), 2.9% acetonitrile (Thermo Scientific) and 0.1% TFA. In addition to individual time points, pooled samples for control and treatment groups were also analysed to investigate differences in the overall secretome. The mass spectrometry proteomics data have been deposited to the ProteomeXchange Consortium via the PRIDE partner repository with the dataset identifier PXD017153 and 10.6019/PXD017153 ^54^. Representative ion chromatograms are shown in Figure S4.

### LC-MS/MS spectra processing and protein identification

Spectral alignment, peak picking, total protein abundance normalisation and peptide/protein quantification were undertaken using Progenesis™ QI 2.0 (Nonlinear Dynamics, Waters). The exported top ten spectra for each feature were then searched against the *Equus caballus* database for peptide and protein identification using PEAKS® Studio 8.5 (Bioinformatics Solutions Inc., Waterloo, Ontario, Canada) software. Search parameters were: precursor mass error tolerance, 10.0 ppm; fragment mass error tolerance, 0.01 Da; precursor mass search type, monoisotopic; enzyme, trypsin; maximum missed cleavages, 1; non-specific cleavage, none; fixed modifications, carbamidomethylation; variable modifications, oxidation or hydroxylation and oxidation (methionine). A filter of a minimum of 2 unique peptides was set for protein identification and quantitation with a false discovery rate (FDR) of 1%.

### 1DSDS PAGE

Media samples for each donor were combined for all time points and analysed via one dimensional sodium dodecyl sulphate polyacrylamide gel electrophoresis (1D SDS PAGE). 1 μg of each sample was added to Laemmli loading buffer Novex™ (Thermo Scientific) producing a final concentration of 15% glycerine, 2.5% SDS, 2.5% Tris (hydroxymethyl) aminomethane, 2.5% HCL and 4% β-mercaptoethanol at pH 6.8 and heated at 95°C for 5 min. Samples were loaded onto a 4-12% Bis-Tris polyacrylamide electrophoresis gel (NuPAGE™ Novex™, Thermo Scientific) and protein separation carried out at 200 V for 30 min at room temperature. Protein bands were visualised via silver staining (Thermo Scientific) following manufacturer instructions. Gel images were converted to 8 bit grey scale and protein band intensities analysed using densitometry with the software Image J (NIH, Bethesda, Maryland).

### Semi-Tryptic Peptide Identification

To identify potential neopeptides a ‘semi-tryptic’ search was undertaken. The same PEAKS® search parameters were used as for protein identification, with the exception that ‘non-specific cleavage’ was altered from ‘none’ to ‘one’. The ‘peptide ion measurements’ file was then exported and analysed using the online neopeptide analyser software tool ^36^.

### Statistical Analysis

Cartilage metabolite profiles were normalised using probabilistic quotient normalisation (PQN) ^55^. Media metabolites were normalised to TSP concentration and protein profiles normalised to total ion current (TIC). Prior to multivariate analysis, metabolite and protein profiles were Pareto scaled ^56^. MetaboAnalyst 3.5 (http://www.metaboanalyst.ca) was used to produce principal component analysis (PCA) plots and provide PC1 (principal component 1) loadings magnitude values. t-tests were carried out using MetaboAnalyst 3.5 (protein and metabolite abundances) and the neopeptide analyser (neopeptides) with p < 0.05 (and a fold change of > 2 for proteins) considered statistically significant. The Benjamini-Hochberg false discovery rate method was applied for correction of multiple testing ^57^. The SPSS 24 software package was used to produce all box plots and PC1 loadings magnitude graphs.

### Pathway Analysis

Pathway analysis was conducted for differentially abundant metabolites using the online tool KEGG for *Equus caballus* ^58^. Differentially abundant proteins (> 2 fold change) were also analysed using KEGG via the STRING database ^59^. Manual cross referencing was conducted of metabolites and proteins found to be significantly different between sample groups to biological pathways. A filter of a minimum of two metabolites/proteins was set for each sample set for inclusion of the relevant biological pathway.

## Results

### NMR Metabolomics

#### Protease inhibitor cocktail, TNF-α and IL-1β metabolite profiles

Protease inhibitor cocktail was found to have high levels of mannitol and thus this metabolite was removed from all analyses. Within the spectral profiles of TNF-α and IL-1β acquired separately, the metabolites acetate, acetone, ethanol, formate, lactate, methanol and succinate were identified. These metabolites were therefore also removed from further analyses.

#### Analysis of Cartilage Metabolites

Acetonitrile metabolite extraction was identified to be highly reproducible with technical replicates clustering within a PCA plot for three separate macroscopically normal cartilage samples (Figure 1). In total 35 metabolites were identified within equine cartilage (Table 1). Of these, following the removal of metabolites previously mentioned, eight were identified as being differentially abundant between control and treatment groups (Figure 2). Glucose and lysine levels were elevated following TNF-α/IL-1β treatment whilst adenosine, alanine, betaine, creatine, myo-inositol and uridine levels decreased. PCA identified that metabolite profiles separated into two distinct clusters, separating control and treatment groups (Figure 2a). PLS-DA (supervised multivariate discriminant analysis) produced a model comprising one component with a good predictive power (R^2^ = 0.89, Q^2^ = 0.82) (Figure 3b). Of the top 25 PC1 loadings, myo-inositol was found to be the most influential cartilage metabolite in separating control and treated samples, followed by glucose, betaine and alanine (Figure 3c).

**Figure 1.**
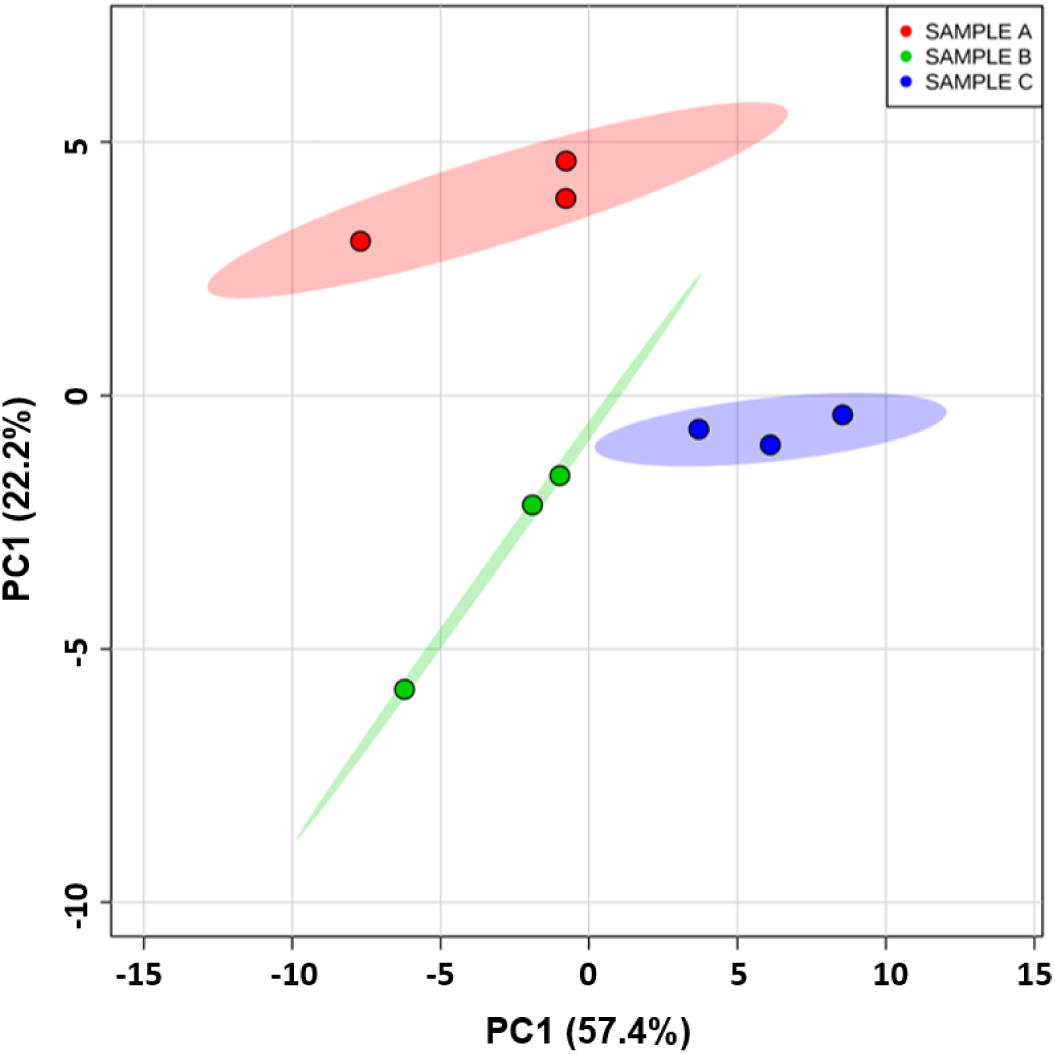
Principal component analysis scores plot identifying high reproducibility of acetonitrile cartilage metabolite extraction (three separate equine donors, technical triplicate for each donor) using 1D ^1^H NMR metabolome analysis. Shaded regions depict 95% confidence regions.

**Figure 2.**
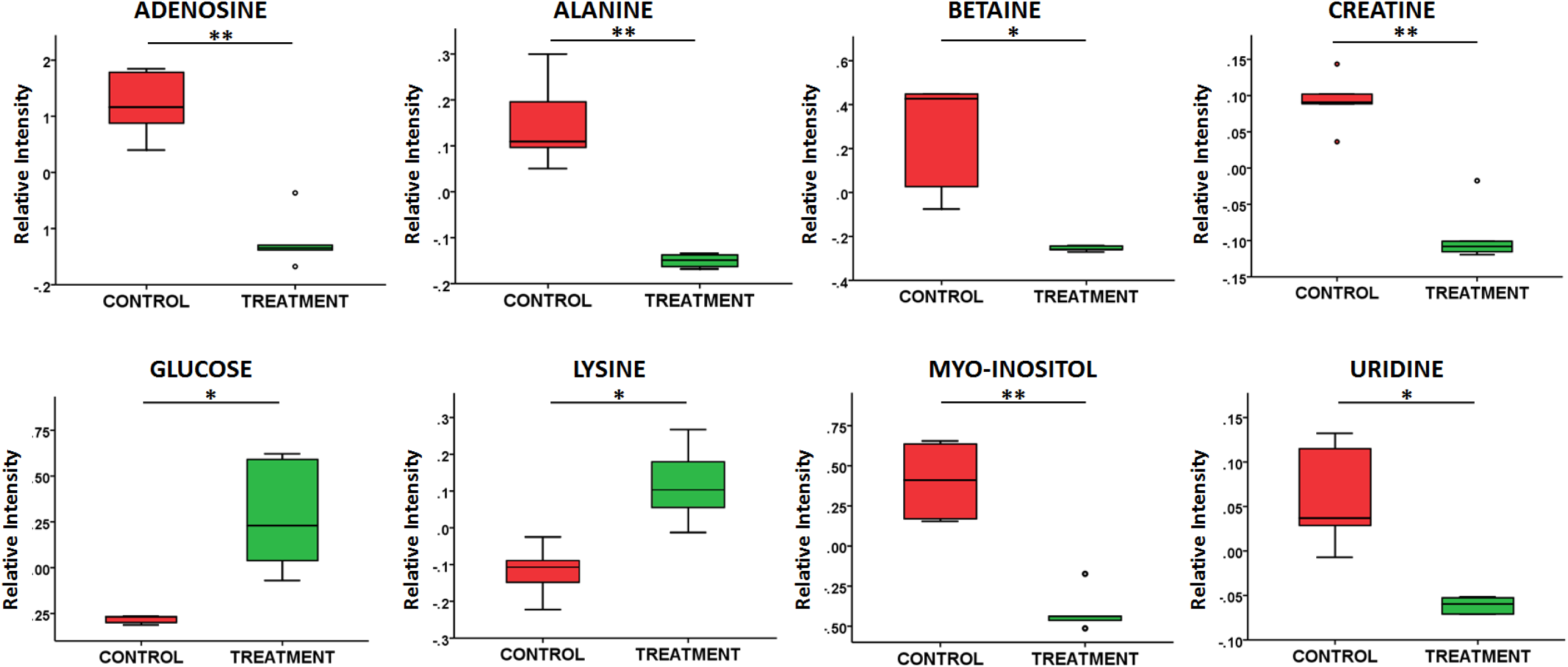
Boxplots of differentially abundant extracted *ex-vivo* equine cartilage metabolites for control (n=5) and following TNF-α/IL-1β treatment (n=5), shown as relative intensities. t-test: * = p < 0.05 and ** = p < 0.01.

**Figure 3.**
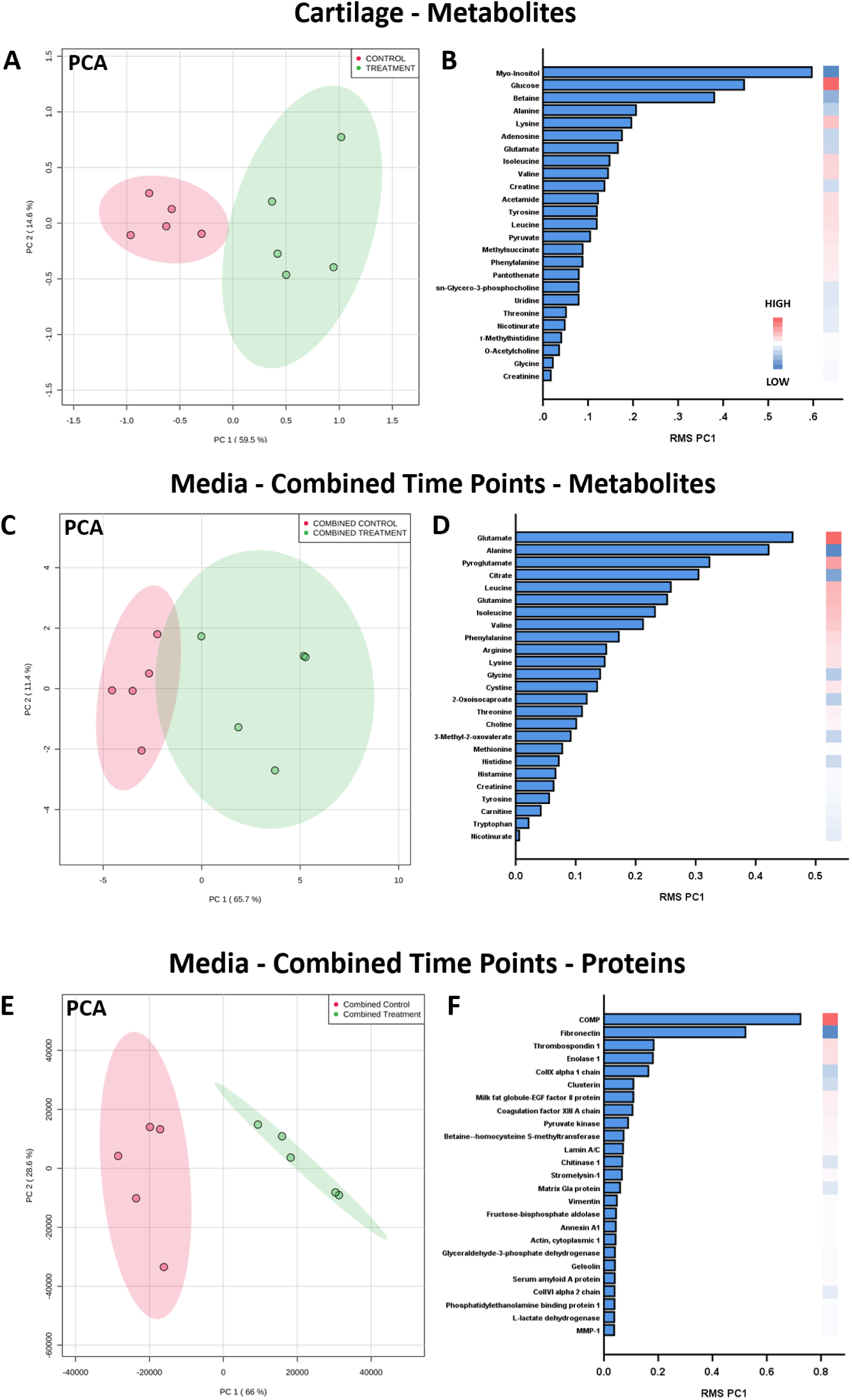
PCA (A, C and E), and PC1 RMS (Principal component 1 root mean square) values (B, D and F) for the 25 components with the highest magnitude for metabolites and proteins present in *ex-vivo* equine cartilage and culture media for combined time points over 8 days, comparing controls (red, n=5) to TNF-α/IL-1β treatment (green, n=5). RMS: High = high in treatment with respect to control, Low = low in treatment with respect to control.

**Table 1.** Metabolites annotated within cartilage and culture media using Chenomx. Metabolites additionally identified using a 1D ^1^H NMR in-house library have been assigned to Metabolomics Standards Initiative (MSI) level 1 ^60^.

| Database Identifier | Metabolite Identification | Cartilage | Cartilage Reliability | Media | Media Reliability |
| --- | --- | --- | --- | --- | --- |
| HMDB00695 | 2-Oxoisocaproate |  |  | Y | MS Level 2 |
| HMDB00491 | 3-Methyl-2-oxovalerate |  |  | Y | MS Level 2 |
| HMDB31645 | Acetamide | Y | MS Level 2 |  |  |
| HMDB00042 | Acetate * | Y | MS Level 1 | Y | MS Level 1 |
| HMDB01659 | Acetone * |  |  | Y | MS Level 2 |
| HMDB00050 | Adenosine | Y | MS Level 2 |  |  |
| HMDB00517 | Arginine |  |  | Y | MS Level 2 |
| HMDB00191 | Aspartate | Y | MS Level 1 |  |  |
| HMDB00043 | Betaine | Y | MS Level 1 |  |  |
| HMDB00097 | Choline |  |  | Y | MS Level 1 |
| HMDB00094 | Citrate | Y | MS Level 1 | Y | MS Level 1 |
| HMDB00064 | Creatine | Y | MS Level 1 |  |  |
| HMDB00562 | Creatinine | Y | MS Level 1 | Y | MS Level 1 |
| HMDB00192 | Cystine |  |  | Y | MS Level 2 |
| HMDB00122 | D-Glucose | Y | MS Level 1 |  |  |
| HMDB04983 | Dimethyl sulfone | Y | MS Level 2 |  |  |
| HMDB00108 | Ethanol * |  |  | Y | MS Level 1 |
| HMDB00142 | Formate * | Y | MS Level 2 | Y | MS Level 2 |
| HMDB00123 | Glycine | Y | MS Level 1 | Y | MS Level 1 |
| HMDB00870 | Histamine |  |  | Y | MS Level 2 |
| HMDB00172 | Isoleucine | Y | MS Level 1 | Y | MS Level 1 |
| HMDB00863 | Isopropanol * |  |  | Y | MS Level 2 |
| HMDB00190 | Lactate * | Y | MS Level 1 | Y | MS Level 1 |
| HMDB00161 | L-Alanine | Y | MS Level 1 | Y | MS Level 1 |
| HMDB00062 | L-Carnitine |  |  | Y | MS Level 2 |
| HMDB00148 | L-Glutamate | Y | MS Level 1 | Y | MS Level 1 |
| HMDB00641 | L-Glutamine | Y | MS Level 1 | Y | MS Level 1 |
| HMDB00177 | L-Histidine |  |  | Y | MS Level 1 |
| HMDB00687 | L-Leucine | Y | MS Level 1 | Y | MS Level 1 |
| HMDB00159 | L-Phenylalanine | Y | MS Level 1 |  |  |
| HMDB00167 | L-Threonine | Y | MS Level 2 | Y | MS Level 2 |
| HMDB00158 | L-Tyrosine | Y | MS Level 1 | Y | MS Level 1 |
| HMDB00883 | L-Valine | Y | MS Level 1 | Y | MS Level 1 |
| HMDB00182 | Lysine | Y | MS Level 1 | Y | MS Level 1 |
| HMDB00765 | Mannitol * |  |  | Y | MS Level 1 |
| HMDB01875 | Methanol * |  |  | Y | MS Level 2 |
| HMDB00696 | Methionine | Y | MS Level 1 | Y | MS Level 1 |
| HMDB01844 | Methylsuccinate | Y | MS Level 2 |  |  |
| HMDB00211 | myo-Inositol | Y | MS Level 1 |  |  |
| HMDB03269 | Nicotinurate | Y | MS Level 2 | Y | MS Level 2 |
| HMDB00895 | O-Acetylcholine | Y | MS Level 2 |  |  |
| HMDB00210 | Pantothenate | Y | MS Level 2 | Y | MS Level 2 |
| HMDB00267 | Pyroglutamate |  |  | Y | MS Level 2 |
| HMDB00243 | Pyruvate | Y | MS Level 1 |  |  |
| HMDB00086 | sn-Glycero-3-phosphocholine | Y | MS Level 2 |  |  |
| HMDB00254 | Succinate * | Y | MS Level 1 | Y | MS Level 1 |
| HMDB00929 | Tryptophan |  |  | Y | MS Level 1 |
| HMDB00300 | Uracil | Y | MS Level 2 |  |  |
| HMDB00296 | Uridine | Y | MS Level 1 |  |  |
| HMDB00001 | τ-Methylhistidine | Y | MS Level 2 | | |

*Y = Yes; \* = Metabolite removed from subsequent analyses*

#### Analysis of Media Metabolites

Spectral quality control via metabolomics standards initiative identified two samples that failed due to salt precipitation and as such were removed from further analyses ^52, 61^. Following metabolite identification and quantification, one sample was identified as an outlier and subsequently removed from statistical analyses. Isopropanol was identified within all media samples. As this was considered a likely contaminant during cartilage culture, together with metabolites previously mentioned, isopropanol was also removed from all analyses. In total, 34 metabolites were identified within the culture media (Table 1). Time points were analysed separately with four, four and six metabolites identified as being differentially abundant between control and treatment groups for 1-2 day, 3-5 day and 6-8 day time points respectively (Figure 4). Choline levels were increased in treated samples compared to controls for all three time points whilst alanine and citrate levels decreased. At 3-5 days glutamate levels were reduced following treatment. At 6-8 days, following treatment, arginine and isoleucine levels were elevated whilst 2-oxoisocaproate and 3-methyl-2-oxovalerate levels were found to decrease. PCA of combined and separated time points identified clear separation between the metabolite profiles of control and treated media samples (Figure 3d and Figure 5a, b and c). Metabolite loadings for PC1 indicate that this separation is driven primarily by alanine and glutamate (Figure 3e and f).

**Figure 4.**
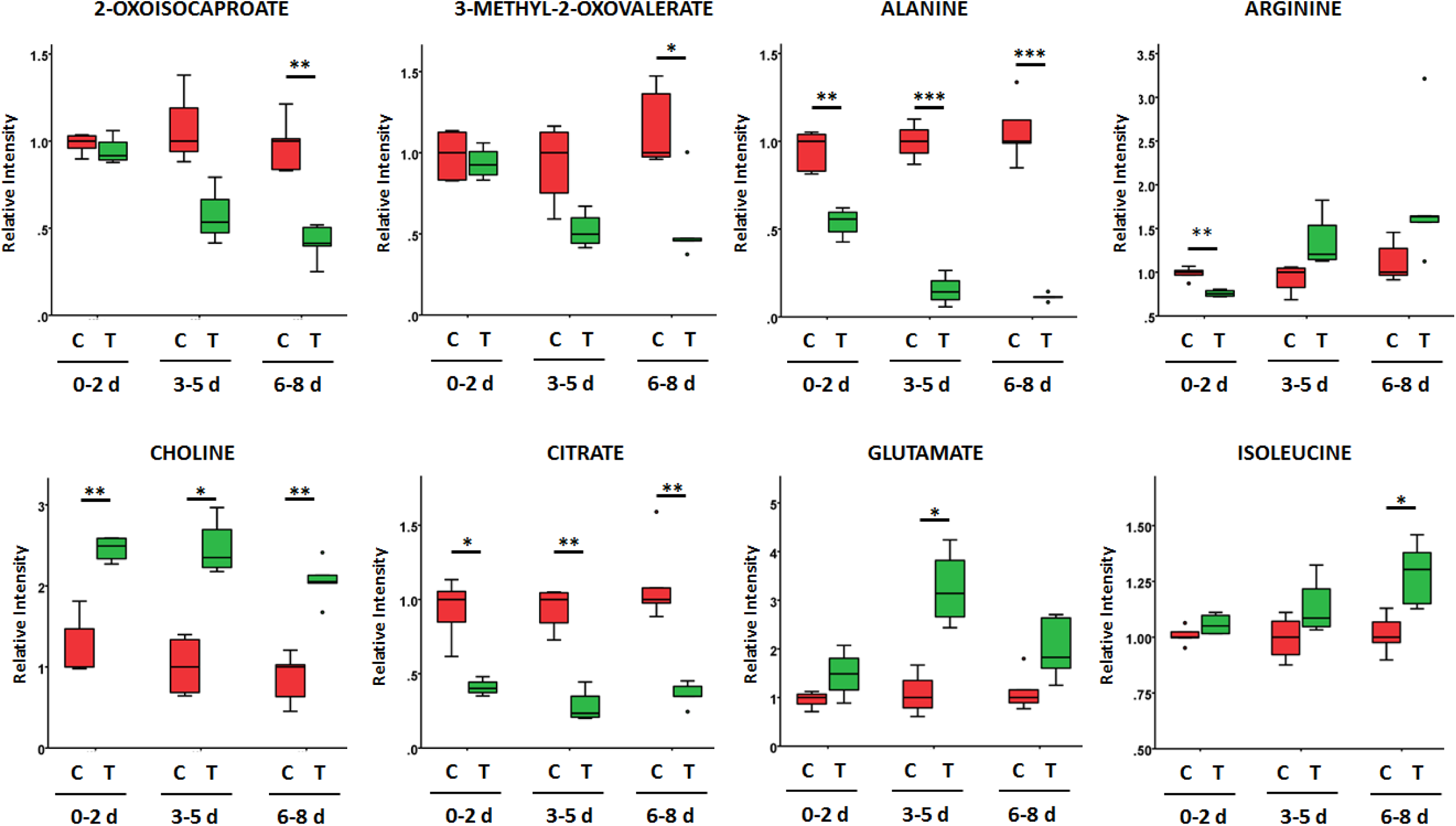
Boxplots of differentially abundant metabolites within the culture media following incubation of *ex-vivo* equine cartilage for control samples (C, red) and following TNF-α/IL-1β treatment (T, green), at 0-2, 3-5 and 6-8 days (d). Metabolite abundances shown as relative intensities. t-test: * = p < 0.05, ** = p < 0.01 and *** = p < 0.001. Control (n=5) and TNF-α/IL-1β treatment (n=5) for each separate time point.

**Figure 5.**
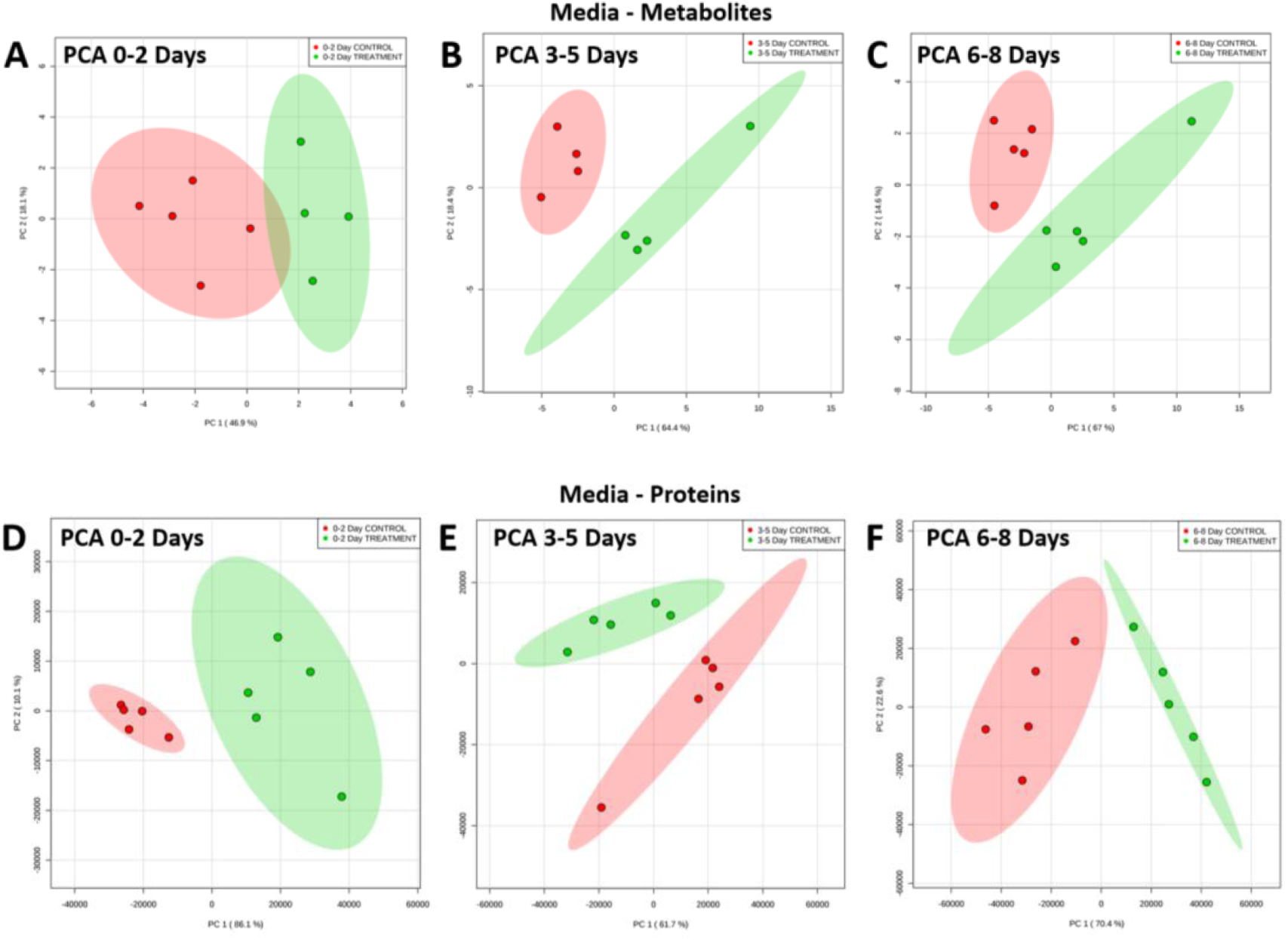
Principal component analysis (PCA) plots of media metabolite (A-C) and protein (D-F) profiles at 0-2, 3-5 and 6-8 days for controls (green, n=5) and TNF-α/IL-1β treatment (red, n=5) of *ex-vivo* equine cartilage.

### LC-MS/MS Proteomics

#### Analysis of Media Proteins

In total, 352 proteins were identified within analysed culture media samples with an elevated number of identified proteins within treated compared to control samples (Figure S5 and Table S1). Of all identified proteins, 131 were identified within TNF-α/IL-1β treated media samples only. When time points were analysed separately, 154, 138 and 72 proteins were identified as being differentially abundant, with > 2 fold change, between control and treatment groups for 1-2 day, 3-5 day and 6-8 day time points respectively.

Supervised PLS-DA analysis of combined protein profiles produced an excellent predictive model (R^2^ = 0.997, Q^2^ = 0.944) with separation primarily driven through elevated COMP and decreased fibronectin following treatment (Figure 3h and 3i). Unsupervised PCA multivariate analysis identified clear discrimination between control and treatment groups at all three time points (Figure 5). At each separated time point the PC1 loadings with the 25 greatest magnitudes corresponding to individual proteins were identified (Figure S6). Box plots in Figure 6 represent proteins which were found to be represented within the top 25 PC1 loadings magnitudes at all three time points (coagulation factor XIII A chain, COMP, enolase 1, Lamin A/C and MMP-3) and extracellular matrix related proteins of interest represented at 2/3 time points (collagen type VI α 2 chain, collagen type X α 1 chain, fibromodulin, fibronectin, matrix Gla protein, MMP1 and vimentin). Coagulation factor XIII A chain, enolase 1 and lamin A/C were elevated at all three time points following treatment. MMP-1 and MMP-3 levels were found to be statistically elevated at 0-2 days only. Fibromodulin and vimentin levels were increased following treatment at both 0-2 days and 3-5 day time points whilst COMP levels increased at 0-2 days and 6-8 days. Collagen type VI α 2 chain and matrix Gla protein levels decreased following treatment at 3-5 days and 6-8 days whilst collagen type X α 1 chain and fibronectin levels statistically decreased at 6-8 days alone.

**Figure 6.**
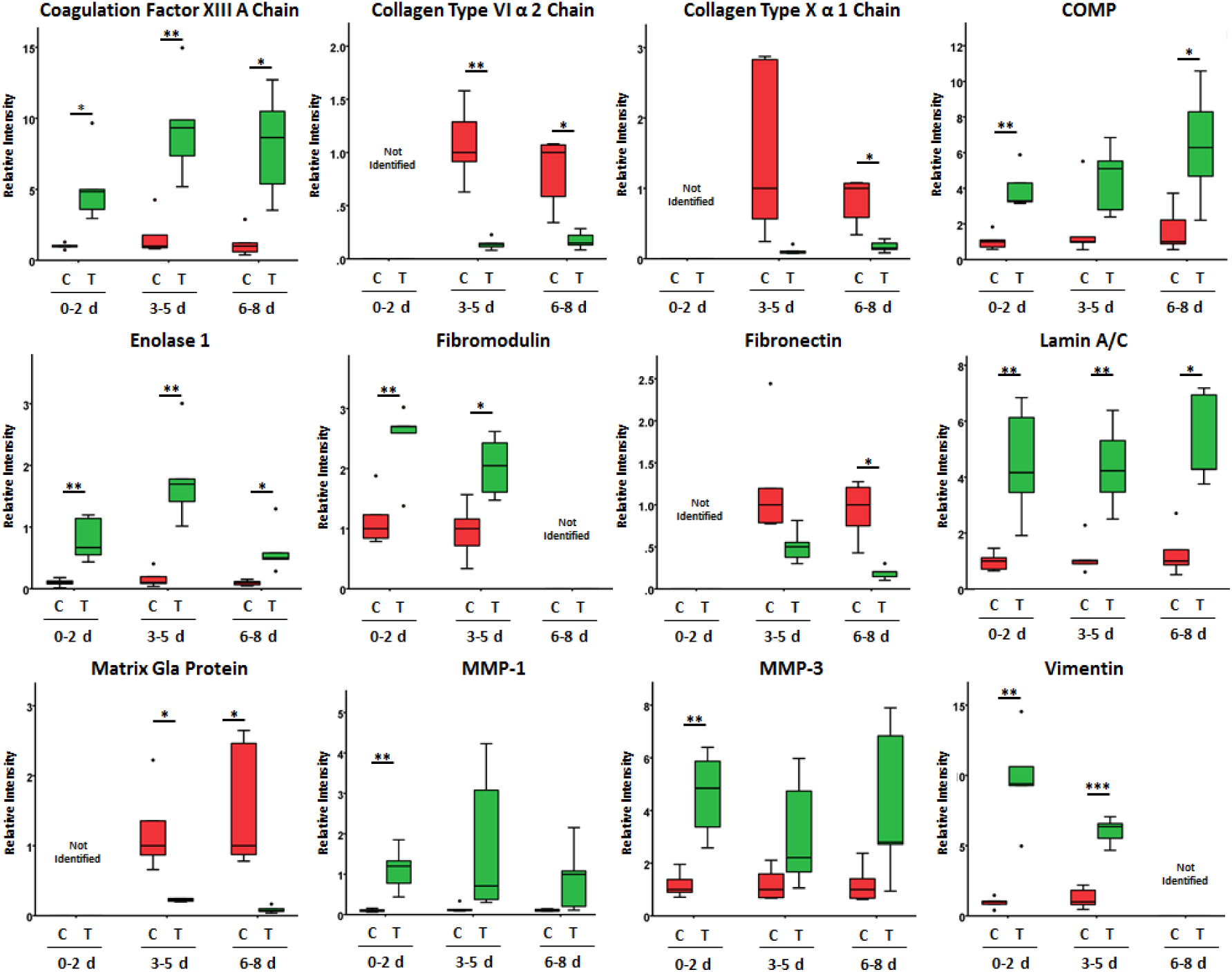
Boxplots of differentially abundant proteins within the culture media following incubation of *ex-vivo* equine cartilage for control samples (C, red) and following TNF-α/IL-1β treatment (T, green), at 0-2, 3-5 and 6-8 days (d). Protein abundances shown as relative intensities. t-test: * = p < 0.05, ** = p < 0.01 and *** = p < 0.001. Control (n=5) and TNF-α/IL-1β treatment (n=5) for each separate time point.

Silver stain analysis of the media profiles for combined time points identified two protein bands which were decreased in abundance following TNF-α/IL-1β treatment, with molecular weights of 160-260 kDa and 260 kDa (Figure 7).

**Figure 7.**
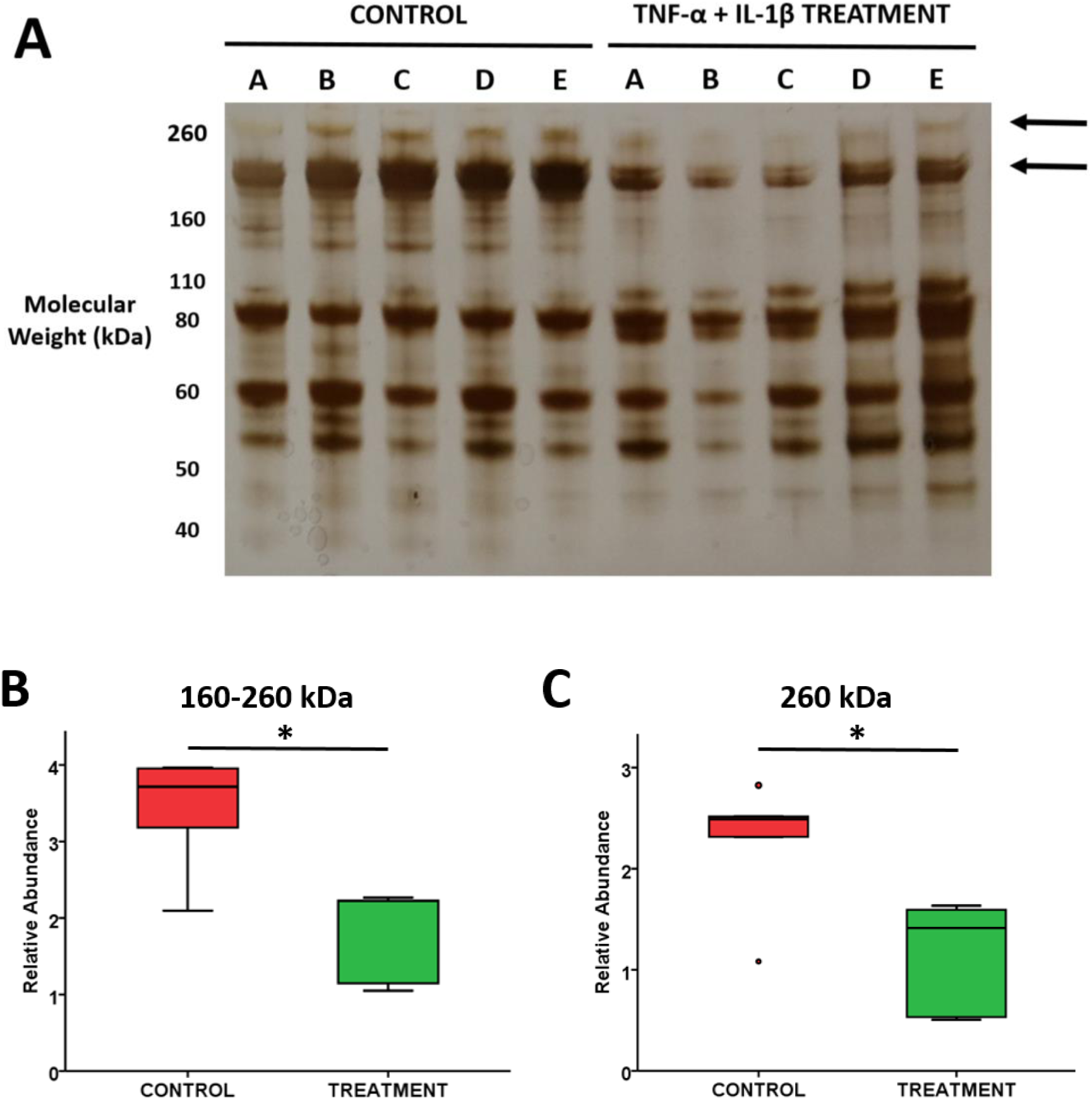
(A) Silver stain identifying media protein profiles (combined for all time points) following incubation of *ex-vivo* equine cartilage for control and TNF-α/IL-1β treated samples. Arrows indicate differentially secreted proteins at approximately 160-260 kDa and 260 kDa. Relative abundances of bands at (B) 160-260 kDa and (C) 260 kDa calculated using densitometry, n=5/group. t-test: * = p < 0.05. A full protein gel image is available in Figure S7.

### Semi-Tryptic Peptides

PCA of all identified semi-tryptic peptides within combined control and combined treated samples identified far less variation within the treatment group (Figure 8). This was also identified for all time points analysed individually (Figure S8). In total, nine potential novel OA neopeptides were identified which were elevated in treated media samples (Table 2). These included semi-tryptic peptides of extracellular matrix proteins aggrecan, cartilage intermediate layer protein, collagen type VI α 2 chain and vimentin.

**Figure 8.**
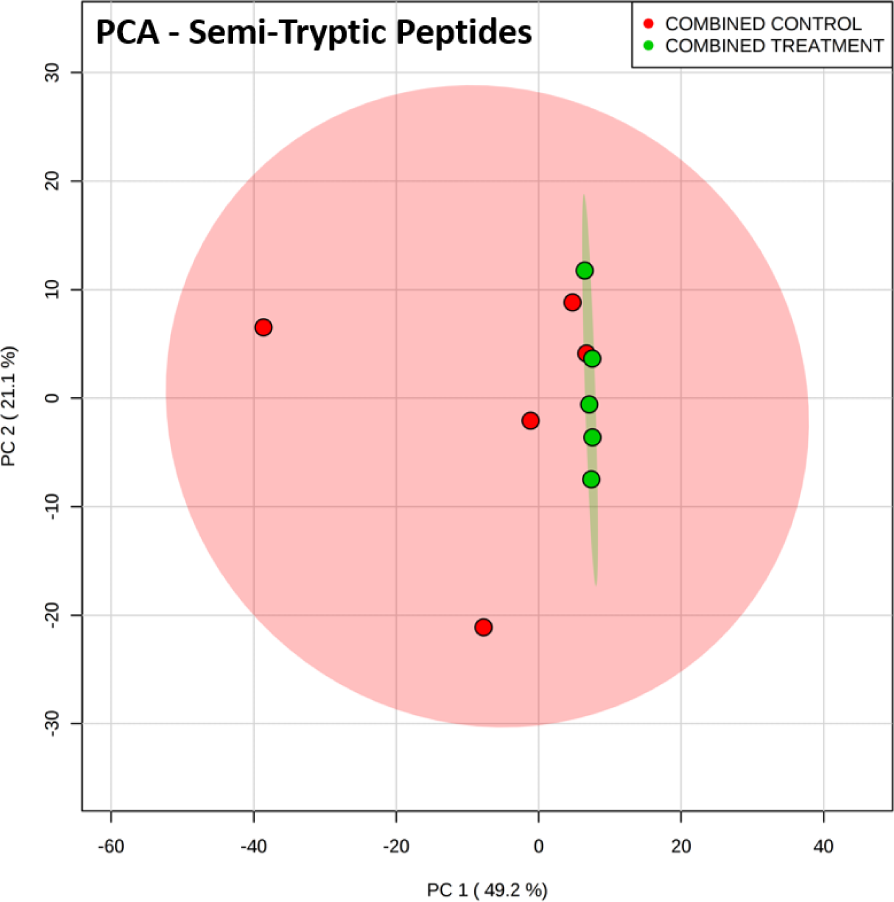
Principal component analysis (PCA) of semi-tryptic peptide profiles within culture media of control (red, n=5) and TNF-α/IL-1β treated (green, n=5) *ex-vivo* equine cartilage. Time points pooled for each individual donor.

**Table 2.**
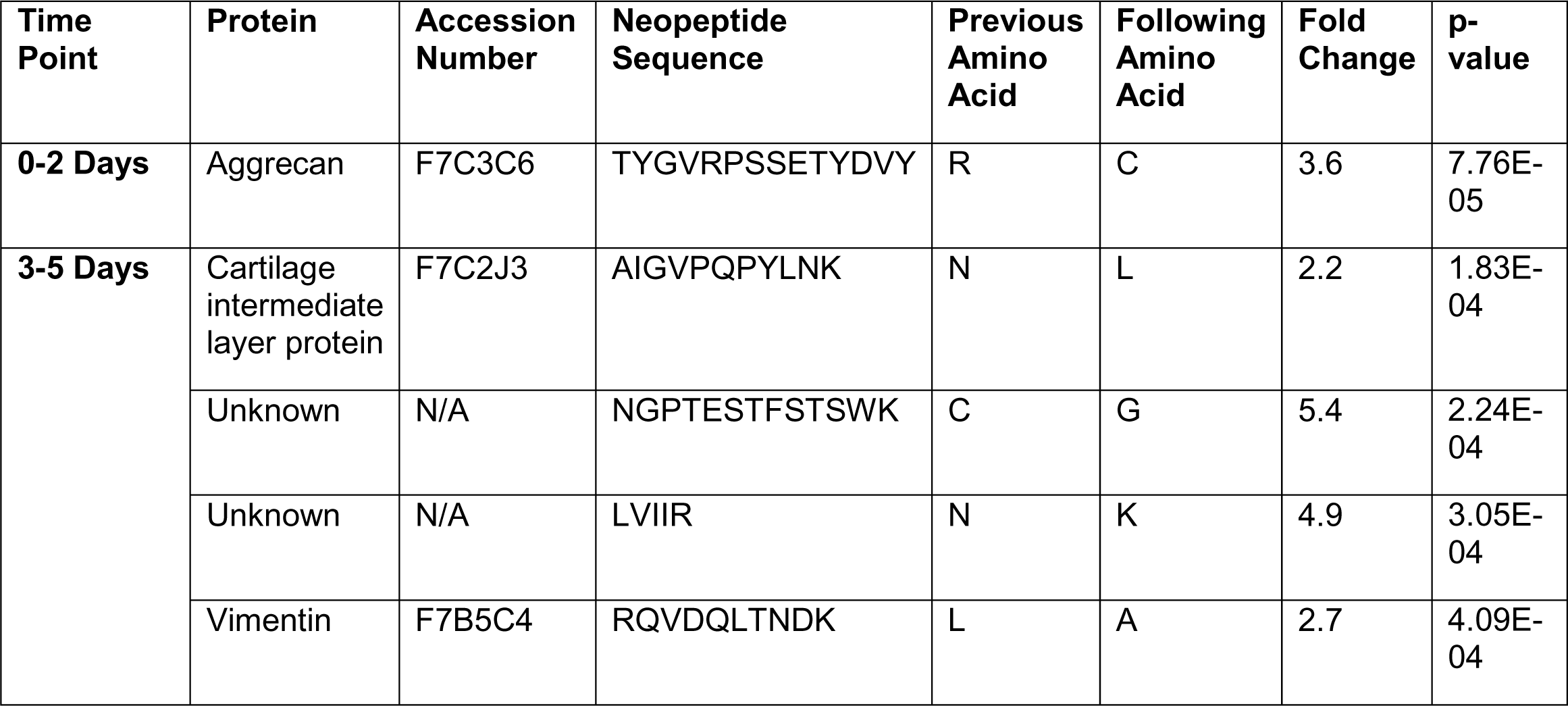
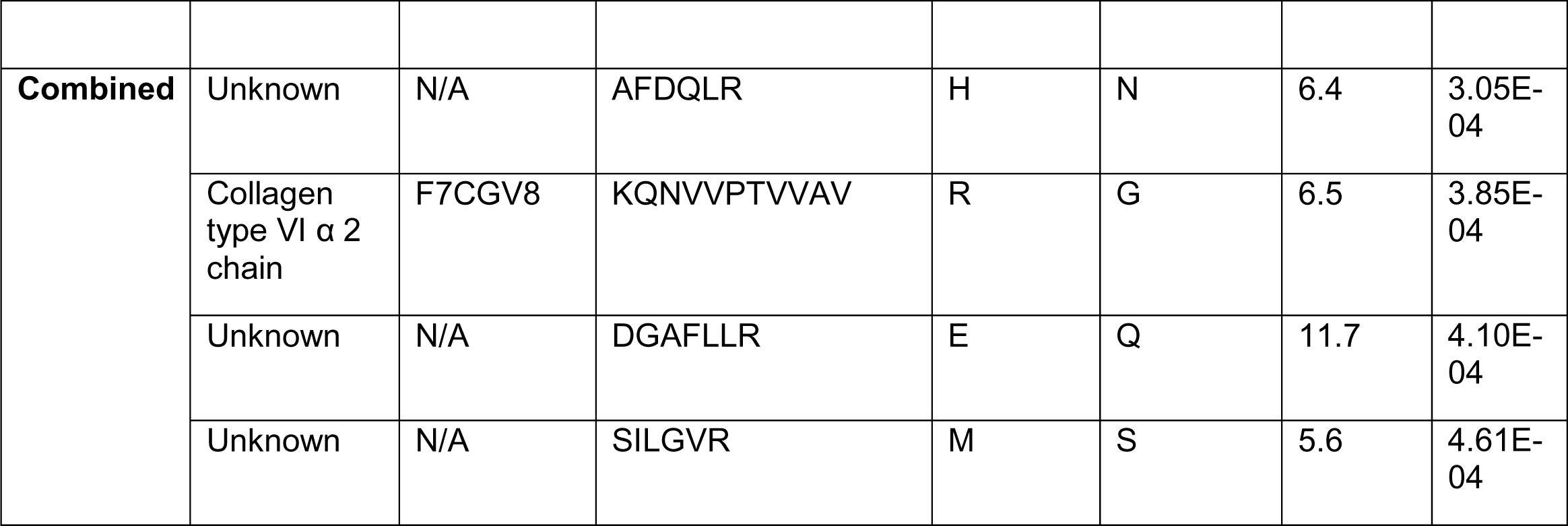
Potential Osteoarthritis Neopeptides. Semi-tryptic peptides of extracellular matrix related and unknown proteins, identified within culture media, with an increased abundance following TNF-α/IL-1β treatment of *ex-vivo* equine cartilage.

| Time Point | Protein | Accession Number | Neopeptide Sequence | Previous Amino Acid | Following Amino Acid | Fold Change | p-value |
| --- | --- | --- | --- | --- | --- | --- | --- |
| 0-2 Days | Aggrecan | F7C3C6 | TYGVRPSSETYDVY | R | C | 3.6 | 7.76E-05 |
| 3-5 Days | Cartilage intermediate layer protein | F7C2J3 | AIGVPQPYLNK | N | L | 2.2 | 1.83E-04 |
|  | Unknown | N/A | NGPTESTFSTSWK | C | G | 5.4 | 2.24E-04 |
|  | Unknown | N/A | LVIIR | N | K | 4.9 | 3.05E-04 |
|  | Vimentin | F7B5C4 | RQVDQLTNDK | L | A | 2.7 | 4.09E-04 |
| **Combined** | Unknown | N/A | AFDQLR | H | N | 6.4 | 3.05E-04 |
| | Collagen type VI $\alpha$ 2 chain | F7CGV8 | KQNVVPTVVAV | R | G | 6.5 | 3.85E-04 |
|  | Unknown | N/A | DGAFLLR | E | Q | 11.7 | 4.10E-04 |
|  | Unknown | N/A | SILGVR | M | S | 5.6 | 4.61E-04 |

### Pathway Analysis

In total, 36 biological pathways were identified which are potentially involved within OA pathogenesis (Table 3). The ‘biosynthesis of amino acids’ and ‘carbon metabolism’ pathways were both highly represented by metabolites and proteins across all three time points. Additionally, ‘ABC Transporters’, ‘alanine, aspartate and glutamate metabolism’ and ‘aminoacyl-tRNA biosynthesis’ were well represented by metabolites across all time points and ‘fructose and mannose metabolism’ and ‘glycolysis/gluconeogenesis’ by proteins across all time points.

**Table 3.** Pathway analysis for differentially abundant metabolites and proteins identified in cartilage and culture media following TNF-α/IL-1β treatment of *ex-vivo* equine cartilage.

|  |  | METABOLITES |  |  |  | PROTEINS |  |  |
| --- | --- | --- | --- | --- | --- | --- | --- | --- |
| Pathway ID | Pathway Description | DA Cartilage Metabolites | DA Media Metabolites |  |  | DA Media Proteins |  |  |
|  |  |  | (0-2 Days) | (3-5 Days) | (6-8 Days) | (0-2 Days) | (3-5 Days) | (6-8 Days) |
| 1210 | 2-Oxocarboxylic acid metabolism |  |  | 2 | 3 |  |  |  |
| 2010 | ABC Transporters | 5 | 3 | 3 | 3 |  |  |  |
| 250 | Alanine, aspartate and glutamate metabolism |  | 2 | 3 | 2 | 4 |  |  |
| 520 | Amino sugar and nucleotide sugar metabolism |  |  |  |  | 6 |  |  |
| 970 | Aminoacyl-tRNA biosynthesis | 2 | 2 | 2 | 2 |  |  |  |
| 4612 | Antigen processing and presentation |  |  |  |  | 5 |  |  |
| 330 | Arginine and proline metabolism |  |  |  |  | 4 |  |  |
| 1230 | Biosynthesis of amino acids | 2 | 3 | 3 | 4 | 13 | 13 | 9 |
| 1200 | Carbon metabolism |  | 2 | 3 | 2 | 13 | 12 | 7 |
| 4610 | Complement and coagulation cascades |  |  |  |  |  | 6 | 6 |
| 270 | Cysteine and methionine metabolism |  |  |  |  | 4 | 4 |  |
| 4512 | ECM-receptor interaction |  |  |  |  |  | 7 |  |
| 4510 | Focal adhesion |  |  |  |  | 7 | 9 |  |
| 51 | Fructose and mannose metabolism |  |  |  |  | 5 | 4 | 3 |
| 52 | Galactose metabolism | 2 |  |  |  | 3 |  |  |
| 480 | Glutathione metabolism |  |  |  |  | 5 | 4 |  |
| 260 | Glycine, serine and threonine metabolism | 2 |  |  |  | 4 | 4 |  |
| 10 | Glycolysis / Gluconeogenesis |  |  |  |  | 12 | 11 | 8 |
| 630 | Glyoxylate and dicarboxylate metabolism |  |  | 2 |  |  |  |  |
| 4978 | Mineral absorption | 2 |  |  | 2 |  |  |  |
| 670 | One carbon pool by folate |  |  |  |  | 4 |  |  |
| 40 | Pentose and glucuronate interconversions |  |  |  |  | 5 |  |  |
| 30 | Pentose phosphate pathway |  |  |  |  | 7 | 7 | 3 |
| 4151 | PI3K-Akt signaling pathway |  |  |  |  |  | 9 |  |
| 640 | Propanoate metabolism |  |  |  |  | 3 | 3 |  |
| 4974 | Protein digestion and absorption | 2 | 2 | 2 | 2 |  | 5 | 5 |
| 4141 | Protein processing in endoplasmic reticulum |  |  |  |  | 8 |  |  |
| 5205 | Proteoglycans in cancer |  |  |  |  | 7 |  |  |
| 230 | Purine metabolism |  |  |  |  | 12 | 7 |  |
| 620 | Pyruvate metabolism |  |  |  |  | 5 | 5 |  |
| 4810 | Regulation of actin cytoskeleton |  |  |  |  | 9 | 7 |  |
| 5150 | Staphylococcus aureus infection |  |  |  |  |  | 4 | 4 |
| 500 | Starch and sucrose metabolism |  |  |  |  | 5 |  |  |
| 430 | Taurine and hypotaurine metabolism |  |  | 2 |  |  |  |  |
| 290 | Valine, leucine and isoleucine biosynthesis |  |  |  | 2 |  |  |  |
| 280 | Valine, leucine and isoleucine degradation |  |  |  | 2 |  |  |  |

*DA = differentially abundant*

## Discussion

In this study, TNF-α/IL-1β treatment of *ex-vivo* equine cartilage explants was used to model OA to gain a greater understanding of OA pathogenesis and identify potential OA markers. ^1^H NMR metabolomic and LC-MS/MS proteomic analysis of culture media at 0-2, 3-5 and 6-8 days was undertaken. In addition, ^1^H NMR metabolomic analysis of acetonitrile extracted cartilage metabolites (following 8 days incubation) was also undertaken.

Within culture media, following TNF-α/IL-1β treatment, elevations in endopeptidases MMP-1 and MMP-3 at 0-2 days, with a similar trend at both other time points, were identified as expected ^62^. Elevated MMP-1 activity has previously been identified within equine OA SF with general MMP activity also found to be correlated to severity of cartilage damage ^63, 64^. Also, as previously reported, elevations in the non-collagenous ECM protein COMP were also identified within the TNF-α/IL-1β equine OA model, with COMP considered a marker of cartilage breakdown ^65, 66^. Clinically, elevated COMP levels have been identified within human OA SF, although within equine OA, one study identified reduced levels with COMP levels being unable to stage the disease ^67, 68^. Fibronectin was identified as a key discriminator between control and treatment groups with reduced secreted fibronectin identified within the media following TNF-α/IL-1β treatment. Additionally, a protein band of molecular weight 160-260 kDa was identified as reduced in treated media compared to control samples via 1D SDS PAGE which may be representing fibronectin (250 kDa), although further techniques, i.e. Western blotting or MS/MS analysis of an in-gel tryptic digest, are required to confirm this ^69, 70^. However, elevated levels of fibronectin have previously been identified within OA SF with fibronectin found to localise at sites of cartilage degeneration and subsequently secreted into the ECM by equine chondrocytes ^71, 72^. The reasons for this possible discrepancy in results between this study and previous studies is currently not known and requires further investigation. It may be that within this study fibronectin has undergone post translational modifications following treatment which may not have been identified via the PEAKS^®^ identification algorithm or resulted in ions which subsequently did not ‘fly’ well during MS analysis and thus were subsequently not identified as peptides.

Following TNF-α/IL-1β treatment, elevations of glucose within the cartilage were identified. This is supported by a previous study which demonstrated that TNF-α and IL-1β upregulate glucose transport in chondrocytes through upregulation of glucose transporter (GLUT)1 and GLUT9 mRNA synthesis with increased levels of glycosylated GLUT1 incorporated into the plasma membrane ^73^. This influx in glucose is likely, at least in part, to be due to the increased energy requirement following cytokine stimulation in the production of MMPs and secretion of Il-6, Il-8, hematopoietic colony-stimulating factor and prostaglandin E_2_ ^74^.

Alanine, arginine, choline and citrate were all identified to be differentially abundant within culture media following TNF-α/IL-1β treatment at the earliest time point, 0-2 days. Within this study, arginine levels were initially identified as decreased following treatment at the earliest time point. A recent study of human plasma also identified arginine to be depleted in knee OA ^75^. The authors proposed this is due to an increased activity of the conversion of arginine to ornithine resulting in an imbalance between cartilage repair and degradation. This is supported by a recent learning and network approach of OA associated metabolites in which arginine and ornithine appeared in about 30% and 25% of the generated models studied ^76^. In addition to this, a reduction in arginine may be reflective of an increased production of nitric oxide (L-arginine being converted to NOH-arginine and subsequently L-citrulline and nitric oxide) as identified in human OA cartilage_77,78._

A ^1^H NMR metabolomics study of equine SF also identified elevated levels of choline in OA ^41^. However, elevated levels of alanine and citrate were also identified whilst these were found to be decreased within our study.

Across the whole study, alanine was found to be a central component in discriminating control and treatment groups. Alanine levels were depleted in treated cartilage extracts compared to controls, with PLS-DA identifying alanine as the fourth most important component in discriminating control and treated cartilage samples. Reduced alanine levels were also identified in human OA cartilage using HRMAS NMR spectroscopy ^50^. Within culture media, at all three time points alanine was depleted in treated samples and KEGG pathway analysis revealed the ‘alanine, aspartate and glutamate metabolism’ pathway to be well represented by both metabolites and proteins. Alanine has previously been identified as a key component of the metabolic urinary OA profile of guinea pigs ^79^.

Isoleucine was elevated within the media during the latter stages of the model (6-8 days). Elevated isoleucine levels have previously been reported within SF of a canine OA model and human OA serum ^39, 80^. Borel *et al.* previously identified elevations of peaks within ^1^H HRMAS NMR spectra of OA cartilage which could be attributed to isoleucine ^49^. Thus the elevations seen in isoleucine may be reflective of cartilage collagen breakdown ^80^.

However, within this study, although a higher abundance was recorded for isoleucine in treated compared to control cartilage, this did not reach statistical significance. Furthermore, elevations in glutamate were identified within culture media at 3-5 days, which may be resultant of the catabolism of collagenous proline through proline oxidase ^81^. Reduced levels of collagen type VI α 2 chain and collagen type X α 1 chain were identified at 3-5 and 6-8 days following cytokine treatment. This may reflect a reduction in collagen synthesis which has previously been identified within other collagen types following TNF-α/IL-1β stimulation ^22^. Therefore these results provide evidence of a disruption in collagen homeostasis and suggest that collagens are being degraded within the model sooner than the 14-28 days previously reported within other *ex-vivo* cartilage OA models ^82, 83^. Coagulation factor XIII A chain, enolase 1, fibromodulin, lamin A/C and vimentin were all identified as being differentially abundant within the culture media at the earliest time point; 0-2 days.

Coagulation factor XIII is a heterotetrameric protein complex which crosslinks fibrin polymers through covalent bonds ^84^. Coagulation factor XIII A chain immunostaining was previously found to be elevated within human articular knee cartilage following IL-1β stimulation ^85^. Sanchez *et al.* identified increased expression of coagulation factor XIII A chain in osteoblasts within the sclerotic zone of OA subchondral bone ^86^. Clinically, remodelling of the subchondral bone is likely to be closely related to cartilage degradation^87^. Within hypertrophic chondrocytes, Nurminkaya *et al.* concluded that cell death and lysis were responsible for the externalisation of the protein (Nurminkaya *et al.*, 1998). However, coagulation factor XIII A chain has also been identified within articular cartilage vesicles, although the underlying externalisation mechanism remains unknown ^89^.

Enolase 1 is a multifunctional glycolytic enzyme which has previously been shown to have increased abundance within an equine articular cartilage model stimulated with IL-1β, as well as increased expression on the cell surface of immune cells during rheumatoid arthritis (RA) ^70, 90^. Lee *et al.* identified apolipoprotein B within RA SF to be a specific ligand to enolase 1, provoking an inflammatory response. Elevated levels of apolipoprotein B have also been associated with human knee OA ^91^. Thus lipid metabolism may operate through this mechanism to regulate chronic inflammation in OA as well as RA ^90^.

Fibromodulin is a small leucine-rich repeat proteoglycan which interacts with collagen fibrils and influences fibrillogenesis rate and fibril structure ^92^. Experimental mice which lack biglycan and fibromodulin have been shown to develop OA in multiple joints ^93, 94^. Neopeptides generated from fibromodulin degradation have also been identified as potential markers of equine articular cartilage degradation ^28^.

Within our study, higher levels of Lamin A/C (intermediate filament protein) were identified within treated media samples ^95^. Lamin A/C has also been identified as being upregulated in human OA cartilage and elevated levels have been implicated in dysregulation of chondrocyte autophagy in ageing and OA ^96, 97^. Thus, our results support these studies, with chondrocyte autophagy targeting a potential novel therapeutic route.

Vimentin is a multifunctional intermediate filament protein ^98^. Within chondrocytes it has been demonstrated that vimentin is likely to be involved in mechanotransduction ^99^. Our results are supported by a previous study which identified elevated levels of cleaved vimentin within human OA cartilage with distortion of the vimentin network evident ^100^.

Due to an elevation in enzymatic activity and breakdown of cartilage during OA, potential biomarkers include ECM degradation fragments ^101^. PCA identified that the semi-tryptic peptide profiles generated from treated equine cartilage was less variable than that of the controls, demonstrating the TNF-α/IL-1β treatment is driving the semi-tryptic peptide profile within the model. Within this study we have identified several semi-tryptic peptides (potential neopeptides) which were identified as being elevated following treatment compared to controls, including degradation products from ECM proteins aggrecan, cartilage intermediate layer protein, vimentin and collagen type VI. None of these potential neopeptides have previously been identified within the literature ^28, 33, 34, 70^.

### Study Limitations

The cytokine preparations used within this study contained various metabolites which, following their removal from subsequent statistical analysis, prevented the analysis of some various metabolites within the experiment. Additionally, culture media was supplemented with a protease inhibitor cocktail at collection in order to inhibit general protein degradation prior to MS proteomic analysis, with results therefore representing the peptide/protein composition during experimentation. However, the high mannitol content prevented analysis of this metabolite within the samples/spectral region. Therefore, when in future using cytokines/supplements for NMR metabolomics, analysing the spectra of different manufacturers/preparations prior to experimentation may be beneficial to identify their associated metabolite profiles, selecting the most appropriate products in order to maximise downstream interpretation of results.

### Further Work

Additional MS based metabolomics analysis of the culture media within this study may be beneficial as NMR and MS are complementary techniques and would therefore expand the number of identified/quantified metabolites, additionally identifying potential lipid and carbohydrate profiles of interest ^9, 38, 43, 102^. In order to confirm the differentially abundant proteins within this study, validation using an orthologous technique e.g. western blotting or enzyme-linked immunosorbent assays is required. Further validation of potential neopeptides could also be carried out through multiple reaction monitoring using a triple-quadrupole mass spectrometer ^103^. Following this, development of monoclonal antibodies specific to neopeptides of interest would enable simpler monitoring of neopeptide abundance in *in vitro*, *ex-vivo* and clinical environments ^104^. Following validations, monitoring the differentially abundant metabolites, proteins and neopeptides within this study within longitudinal SF samples from OA horses would identify translation of these findings to a clinical setting and the eventual generation of clinically applicable diagnostic tests.

## Conclusion

In conclusion, this is the first study to use a multi ‘omics’ approach to simultaneously investigate the metabolomic profile of *ex-vivo* cartilage and metabolomic/proteomic profiles of culture media using the TNF-α/IL-1β *ex-vivo* OA cartilage model. We have identified a panel of metabolites and proteins which are differentially abundant within an early phase of the OA model, 0-2 days, which may provide further information on the underlying disease pathogenesis and well as potential to translate to clinical markers. This study has also identified a panel of potential, ECM derived, neopeptides which have potential to help enable OA stratification as well as provide potential novel therapeutic targets.

## Supporting Information

### Liquid Chromatography Tandem Mass Spectrometry - Detailed Methods

**Figure S1.** Five post mortem equine metacarpophalangeal joints used for *ex-vivo* cartilage culture.

**Figure S2.** Experimental design for *ex-vivo* equine cartilage culture +/-TNF-α/IL-1β treatment. MCP = metacarpophalangeal.

**Figure S3.** 1D ^1^H nuclear magnetic resonance spectral quantile plots of cartilage - 8 days in control media, cartilage - 8 days in TNF-α/IL-1β treated media, control media (all time points combined) and TNF-α/IL-1β treated media (all time points combined).

**Figure S4.** Representative culture media ion chromatograms of combined time points for control and TNF-α/IL-1β treated equine *ex-vivo* cartilage explants using a 60 min liquid chromatography gradient.

**Figure S5.** Number of proteins identified within culture media for (A) combined and (B) individual sample time points for controls and TNF-α/IL-1β treated *ex-vivo* equine cartilage. Combined

**Figure S6.** PC1 RMS (Principal component 1 root mean square) values for the 25 components with the highest magnitude for differentially abundant proteins present within culture media at (A) 0-2 days, (B) 3-5 days and (C) 6-8 days following TNF-α/IL-1β treatment of *ex-vivo* equine cartilage. n=5 for each time point. RMS: High = high in treatment with respect to control, Low = low in treatment with respect to control.

**Figure S7.** Full Protein Gel Image used in Figure 7. Silver stain identifying media protein profiles (combined for all time points) following incubation of *ex-vivo* equine cartilage for control and TNF-α/IL-1β treated samples.

**Figure S8.** Principal component analyses of semi-tryptic peptide profiles within culture media of control and TNF-α/IL-1β treated *ex-vivo* equine cartilage at 0-2 days, 3-5 days and 6-8 days.

**Table S1.** Proteins identified within culture media of control and TNF-α/IL-1β treated *ex-vivo* equine cartilage time points.

## Ethics

Cartilage samples were collected as a by-product of the agricultural industry. The Animals (Scientific Procedures) Act 1986, Schedule 2, does not define collection from these sources as scientific procedures and ethical approval was therefore not required.

## Corresponding author

James R Anderson Tel: 01517949287

## Author Contributions

Wrote the manuscript (J.A.), revised the manuscript (J.A., M.M.P., P.C., M.J.P.), collected cartilage samples (J.A.), experimental procedures (J.A., L.F., M.M.P.), analysed the data (J.A., L.F., M.M.P.), experimental design (J.A., M.M.P., P.C., M.J.P.). All authors read and approved the final manuscript.

## Notes

The authors declare no competing financial interest.

## Acknowledgements

The authors would like to thank staff at F Drury and Sons abattoir, Swindon for their assistance in sample collection, members of the Centre for Protein Research, University of Liverpool including Professor Rob Beynon and Dr Philip Brownridge and Mr Jake Ellis, Cardiff University, for undertaking NMR file depositions..

## Abbreviations

ADAMTS: A disintegrin and metalloproteinase with thrombospondin motifs
CPMG: Carr-Purcell-Meiboom-Gill
COMP: Cartilage oligomeric matrix protein
DMEM: Dulbecco’s modified Eagle’s medium
EDTA: Ethylenediaminetetraacetic acid
ECM: Extracellular matrix
FDR: False discovery rate
FCS: Foetal calf serum
GLUT: Glucose transporter
HRMAS: High resolution magical angle spinning
IL-1β: Interleukin-1β
MS: Mass spectrometry
MMP: Matrix metalloproteinase
MSI: Metabolomics Standards Initiative
NMR: Nuclear magnetic resonance
1D SDS PAGE: One dimensional sodium dodecyl sulphate polyacrylamide gel electrophoresis
OA: Osteoarthritis
PBS: Phosphate buffered saline
PCA: Principal component analysis
PC1: Principal component 1
PQN: Probabilistic quotient normalisation
RA: Rheumatoid arthritis
RMS: Root mean square
SF: Synovial fluid
TIC: Total ion current
TFA: Trifluoroacetic acid
TSP: Trimethylsilyl propionate
TNF-α: Tumour necrosis factor-α

## References

1. Truong, L.-H.; Kuliwaba, J. S.; Tsangari, H.; Fazzalari, N. L. Differential Gene Expression of Bone Anabolic Factors and Trabecular Bone Architectural Changes in the Proximal Femoral Shaft of Primary Hip Osteoarthritis Patients. Arthritis Res. Ther. 2006, 8 (6), R188–R188. https://doi.org/10.1186/ar2101.

2. Kramer, C. M.; Tsang, A. S.; Koenig, T.; Jeffcott, L. B.; Dart, C. M.; Dart, A. J. Survey of the Therapeutic Approach and Efficacy of Pentosan Polysulfate for the Prevention and Treatment of Equine Osteoarthritis in Veterinary Practice in Australia. Aust Vet J 2014, 92 (12), 482–487. https://doi.org/10.1111/avj.12266.

3. Ireland, J. L.; Clegg, P. D.; McGowan, C. M.; Platt, L.; Pinchbeck, G. L. Factors Associated with Mortality of Geriatric Horses in the United Kingdom. Prev Vet Med 2011, 101 (3–4), 204–218. https://doi.org/10.1016/j.prevetmed.2011.06.002.

4. Ireland, J. L.; Clegg, P. D.; McGowan, C. M.; McKane, S. A.; Chandler, K. J.; Pinchbeck, G. L. Disease Prevalence in Geriatric Horses in the United Kingdom: Veterinary Clinical Assessment of 200 Cases. Equine Vet J 2012, 44 (1), 101–106. https://doi.org/10.1111/j.2042-3306.2010.00361.x.

5. Caron, J.; Genovese, R. Principles and Practices of Joint Disease Treatment. In Diagnosis and management of lameness in the horse; Ross, M., Dyson, S., Eds.; W.B. Saunders: Philadelphia, 2003; pp 746–764.

6. Struglics, A.; Larsson, S.; Pratta, M. A.; Kumar, S.; Lark, M. W.; Lohmander, L. S. Human Osteoarthritis Synovial Fluid and Joint Cartilage Contain Both Aggrecanase- and Matrix Metalloproteinase-Generated Aggrecan Fragments. Osteoarthr. Cartil. 2006, 14 (2), 101–113. https://doi.org/10.1016/j.joca.2005.07.018.

7. Li, Y.; Xu, L.; Olsen, B. R. Lessons from Genetic Forms of Osteoarthritis for the Pathogenesis of the Disease. Osteoarthr. Cartil. 2007, 15 (10), 1101–1105. https://doi.org/10.1016/j.joca.2007.04.013.

8. Marhardt, K.; Muurahainen, N. Development of a Disease-Modifying OA Drug (DMOAD) in Knee Osteoarthritis: The Example of Sprifermin. Drug Res. (Stuttg). 2015, 65 (S 01), S13–S13. https://doi.org/10.1055/s-0035-1558063.

9. Anderson, J. R.; Phelan, M. M.; Clegg, P. D.; Peffers, M. J.; Rubio-Martinez, L. M. Synovial Fluid Metabolites Differentiate between Septic and Nonseptic Joint Pathologies. J. Proteome Res. 2018, 17 (8), 2735–2743. https://doi.org/10.1021/acs.jproteome.8b00190.

10. Brommer, H.; van Weeren, P. R.; Brama, P. A. New Approach for Quantitative Assessment of Articular Cartilage Degeneration in Horses with Osteoarthritis. Am J Vet Res 2003, 64 (1), 83–87.

11. Hunter, D. J.; Nevitt, M.; Losina, E.; Kraus, V. Biomarkers for Osteoarthritis: Current Position and Steps towards Further Validation. Best Pract. Res. Clin. Rheumatol. 2014, 28 (1), 61–71. https://doi.org/10.1016/j.berh.2014.01.007.

12. McIlwraith, C. W.; Kawcak, C. E.; Frisbie, D. D.; Little, C. B.; Clegg, P. D.; Peffers, M. J.; Karsdal, M. A.; Ekman, S.; Laverty, S.; Slayden, R. A.;, et al. Biomarkers for Equine Joint Injury and Osteoarthritis. J. Orthop. Res. 2018, 36 (3), 823–831. https://doi.org/10.1002/jor.23738.

13. Wojdasiewicz, P.; Poniatowski, Ł. A.; Szukiewicz, D. The Role of Inflammatory and Anti-Inflammatory Cytokines in the Pathogenesis of Osteoarthritis. Mediators Inflamm. 2014, 2014, 561459. https://doi.org/10.1155/2014/561459.

14. Wang, J.; Markova, D.; Anderson, D. G.; Zheng, Z.; Shapiro, I. M.; Risbud, M. V. TNF-α and IL-1β Promote a Disintegrin-like and Metalloprotease with Thrombospondin Type I Motif-5-Mediated Aggrecan Degradation through Syndecan-4 in Intervertebral Disc. J. Biol. Chem. 2011, 286 (46), 39738–39749. https://doi.org/10.1074/jbc.M111.264549.

15. Fernandes, J. C.; Martel-Pelletier, J.; Pelletier, J.-P. The Role of Cytokines in Osteoarthritis Pathophysiology. Biorheology 2002, 39 (1–2), 237–246.

16. Goldring, S. R.; Goldring, M. B. The Role of Cytokines in Cartilage Matrix Degeneration in Osteoarthritis. Clin. Orthop. Relat. Res. 2004, *427S*, 27–36. https://doi.org/10.1097/01.blo.0000144854.66565.8f.

17. Westacott, C. I.; Whicher, J. T.; Barnes, I. C.; Thompson, D.; Swan, A. J.; Dieppe, P. A. Synovial Fluid Concentration of Five Different Cytokines in Rheumatic Diseases. Ann. Rheum. Dis. 1990, 49 (9), 676–681. https://doi.org/10.1136/ARD.49.9.676.

18. Bertuglia, A.; Pagliara, E.; Grego, E.; Ricci, A.; Brkljaca-Bottegaro, N. Pro-Inflammatory Cytokines and Structural Biomarkers Are Effective to Categorize Osteoarthritis Phenotype and Progression in Standardbred Racehorses over Five Years of Racing Career. BMC Vet. Res. 2016, 12 (1), 246. https://doi.org/10.1186/s12917-016-0873-7.

19. Ma, T.-W.; Li, Y.; Wang, G.-Y.; Li, X.-R.; Jiang, R.-L.; Song, X.-P.; Zhang, Z.-H.; Bai, H.; Li, X.; Gao, L. Changes in Synovial Fluid Biomarkers after Experimental Equine Osteoarthritis. *J*. Vet. Res. 2017, 61 (4), 503–508. https://doi.org/10.1515/jvetres-2017-0056.

20. Williams, A. Proteomic Studies of an Explant Model of Equine Articular Cartilage in Response to Proinflammatory and Anti-Inflammatory Stimuli, University of Nottingham, 2014.

21. Stevens, A. L.; Wishnok, J. S.; Chai, D. H.; Grodzinsky, A. J.; Tannenbaum, S. R. A Sodium Dodecyl Sulfate-Polyacrylamide Gel Electrophoresis-Liquid Chromatography Tandem Mass Spectrometry Analysis of Bovine Cartilage Tissue Response to Mechanical Compression Injury and the Inflammatory Cytokines Tumor Necrosis Factor α and Interleukin. Arthritis Rheum. 2008, 58 (2), 489–500. https://doi.org/10.1002/art.23120.

22. Stevens, A. L.; Wishnok, J. S.; White, F. M.; Grodzinsky, A. J.; Tannenbaum, S. R. Mechanical Injury and Cytokines Cause Loss of Cartilage Integrity and Upregulate Proteins Associated with Catabolism, Immunity, Inflammation, and Repair. Mol. Cell. Proteomics 2009, 8 (7), 1475–1489. https://doi.org/10.1074/mcp.M800181-MCP200.

23. Pretzel, D.; Pohlers, D.; Weinert, S.; Kinne, R. W. In Vitro Model for the Analysis of Synovial Fibroblast-Mediated Degradation of Intact Cartilage. Arthritis Res. Ther. 2009, 11 (1), R25. https://doi.org/10.1186/ar2618.

24. De Ceuninck, F.; Dassencourt, L.; Anract, P. The Inflammatory Side of Human Chondrocytes Unveiled by Antibody Microarrays. Biochem. Biophys. Res. Commun. 2004, 323 (3), 960–969. https://doi.org/10.1016/J.BBRC.2004.08.184.

25. Cillero-Pastor, B.; Ruiz-Romero, C.; Caramés, B.; López-Armada, M. J.; Blanco, F. J. Proteomic Analysis by Two-Dimensional Electrophoresis to Identify the Normal Human Chondrocyte Proteome Stimulated by Tumor Necrosis Factor α and Interleukin-1β. Arthritis Rheum. 2010, 62 (3), 802–814. https://doi.org/10.1002/art.27265.

26. Barksby, H. E.; Milner, J. M.; Patterson, A. M.; Peake, N. J.; Hui, W.; Robson, T.; Lakey, R.; Middleton, J.; Cawston, T. E.; Richards, C. D.;, et al. Matrix Metalloproteinase 10 Promotion of Collagenolysis via Procollagenase Activation: Implications for Cartilage Degradation in Arthritis. Arthritis Rheum. 2006, 54 (10), 3244–3253. https://doi.org/10.1002/art.22167.

27. de Hoog, C. L.; Mann, M. Proteomics. Annu. Rev. Genomics Hum. Genet. 2004, 5 (1), 267–293. https://doi.org/10.1146/annurev.genom.4.070802.110305.

28. Peffers, M. J.; Thornton, D. J.; Clegg, P. D. Characterization of Neopeptides in Equine Articular Cartilage Degradation. J. Orthop. Res. 2016, 34 (1), 106–120. https://doi.org/10.1002/jor.22963.

29. Polur, I.; Lee, P. L.; Servais, J. M.; Xu, L.; Li, Y. Role of HTRA1, a Serine Protease, in the Progression of Articular Cartilage Degeneration. Histol. Histopathol. 2010, 25 (5), 599–608. https://doi.org/10.14670/HH-25.599.

30. Ben-Aderet, L.; Merquiol, E.; Fahham, D.; Kumar, A.; Reich, E.; Ben-Nun, Y.; Kandel, L.; Haze, A.; Liebergall, M.; Kosińska, M. K.;, et al. Detecting Cathepsin Activity in Human Osteoarthritis via Activity-Based Probes. Arthritis Res. Ther. 2015, 17 (1), 69. https://doi.org/10.1186/s13075-015-0586-5.

31. Peffers, M. J.; Smagul, A.; Anderson, J. R. Proteomic Analysis of Synovial Fluid: Current and Potential Uses to Improve Clinical Outcomes. Expert Rev. Proteomics 2019, 16 (4), 287–302. https://doi.org/10.1080/14789450.2019.1578214.

32. Miller, R. E.; Ishihara, S.; Tran, P. B.; Golub, S. B.; Last, K.; Miller, R. J.; Fosang, A. J.; Malfait, A.-M. An Aggrecan Fragment Drives Osteoarthritis Pain through Toll-like Receptor 2. JCI Insight 2018, 3 (6), 1–9. https://doi.org/10.1172/JCI.INSIGHT.95704.

33. Peffers, M. J.; Cillero-Pastor, B.; Eijkel, G. B.; Clegg, P. D.; Heeren, R. M. Matrix Assisted Laser Desorption Ionization Mass Spectrometry Imaging Identifies Markers of Ageing and Osteoarthritic Cartilage. Arthritis Res. Ther. 2014, 16 (3), R110. https://doi.org/10.1186/ar4560.

34. Peffers, M. J.; McDermott, B.; Clegg, P. D.; Riggs, C. M. Comprehensive Protein Profiling of Synovial Fluid in Osteoarthritis Following Protein Equalization. Osteoarthr. Cartil. 2015, 23 (7), 1204–1213. https://doi.org/10.1016/j.joca.2015.03.019.

35. Skiöldebrand, E.; Ekman, S.; Mattsson Hultén, L.; Svala, E.; Björkman, K.; Lindahl, A.; Lundqvist, A.; Önnerfjord, P.; Sihlbom, C.; Rüetschi, U. Cartilage Oligomeric Matrix Protein Neoepitope in the Synovial Fluid of Horses with Acute Lameness: A New Biomarker for the Early Stages of Osteoarthritis. Equine Vet. J. 2017, 49 (5), 662–667. https://doi.org/10.1111/evj.12666.

36. Peffers, M.; Jones, A. R.; McCabe, A.; Anderson, J. Neopeptide Analyser: A Software Tool for Neopeptide Discovery in Proteomics Data. Wellcome Open Res. 2017, 2, 24. https://doi.org/10.12688/wellcomeopenres.11275.1.

37. Beckonert, O.; Keun, H. C.; Ebbels, T. M. D.; Bundy, J.; Holmes, E.; Lindon, J. C.; Nicholson, J. K. Metabolic Profiling, Metabolomic and Metabonomic Procedures for NMR Spectroscopy of Urine, Plasma, Serum and Tissue Extracts. Nat. Protoc. 2007, 2 (11), 2692–2703. https://doi.org/10.1038/nprot.2007.376.

38. Beltran, A.; Suarez, M.; Rodríguez, M. A.; Vinaixa, M.; Samino, S.; Arola, L.; Correig, X.; Yanes, O. Assessment of Compatibility between Extraction Methods for NMR- and LC/MS-Based Metabolomics. Anal. Chem. 2012, 84 (14), 5838–5844. https://doi.org/10.1021/ac3005567.

39. Damyanovich, A. Z.; Staples, J. R.; Chan, A. D.; Marshall, K. W. Comparative Study of Normal and Osteoarthritic Canine Synovial Fluid Using 500 MHz 1H Magnetic Resonance Spectroscopy. J Orthop Res 1999, 17 (2), 223–231. https://doi.org/10.1002/jor.1100170211.

40. Hugle, T.; Kovacs, H.; Heijnen, I. A.; Daikeler, T.; Baisch, U.; Hicks, J. M.; Valderrabano, V. Synovial Fluid Metabolomics in Different Forms of Arthritis Assessed by Nuclear Magnetic Resonance Spectroscopy. Clin Exp Rheumatol 2012, 30 (2), 240–245.

41. Lacitignola, L.; Fanizzi, F. P.; Francioso, E.; Crovace, A. 1H NMR Investigation of Normal and Osteo-Arthritic Synovial Fluid in the Horse. Vet. Comp. Orthop. Traumatol. 2008, 21 (1), 85–88. https://doi.org/10.3415/VCOT-06-12-0101.

42. Mickiewicz, B.; Heard, B. J.; Chau, J. K.; Chung, M.; Hart, D. A.; Shrive, N. G.; Frank, C. B.; Vogel, H. J. Metabolic Profiling of Synovial Fluid in a Unilateral Ovine Model of Anterior Cruciate Ligament Reconstruction of the Knee Suggests Biomarkers for Early Osteoarthritis. J. Orthop. Res. 2015, 33 (1), 71–77. https://doi.org/10.1002/jor.22743.

43. Mickiewicz, B.; Kelly, J. J.; Ludwig, T. E.; Weljie, A. M.; Wiley, J. P.; Schmidt, T. A.; Vogel, H. J. Metabolic Analysis of Knee Synovial Fluid as a Potential Diagnostic Approach for Osteoarthritis. J Orthop Res 2015, 33 (11), 1631–1638. https://doi.org/10.1002/jor.22949.

44. Anderson, J. R.; Chokesuwattanaskul, S.; Phelan, M. M.; Welting, T. J. M.; Lian, L.-Y.; Peffers, M. J.; Wright, H. L. 1 H NMR Metabolomics Identifies Underlying Inflammatory Pathology in Osteoarthritis and Rheumatoid Arthritis Synovial Joints. J. Proteome Res. 2018, 17 (11), 3780–3790. https://doi.org/10.1021/acs.jproteome.8b00455.

45. Graham, R. J. T. Y.; Anderson, J. R.; Phelan, M. M.; Cillan-Garcia, E.; Bladon, B. M.; Taylor, S. E. Metabolomic Analysis of Synovial Fluid from Thoroughbred Racehorses Diagnosed with Palmar Osteochondral Disease Using Magnetic Resonance Imaging. Equine Vet. J. 2019, evj.13199. https://doi.org/10.1111/evj.13199.

46. Ling, W.; Regatte, R. R.; Schweitzer, M. E.; Jerschow, A. Characterization of Bovine Patellar Cartilage by NMR. NMR Biomed. 2008, 21 (3), 289–295. https://doi.org/10.1002/nbm.1193.

47. Schiller, J.; Huster, D.; Fuchs, B.; Naji, L.; Kaufmann, J.; Arnold, K. Evaluation of Cartilage Composition and Degradation by High-Resolution Magic-Angle Spinning Nuclear Magnetic Resonance. In Cartilage and Osteoarthritis; Humana Press: New Jersey, 2004; pp 267–286. https://doi.org/10.1385/1-59259-821-8:267.

48. Schiller, J.; Naji, L.; Huster, D.; Kaufmann, J.; Arnold, K. 1H And13C HR-MAS NMR Investigations on Native and Enzymatically Digested Bovine Nasal Cartilage. Magma Magn. Reson. Mater. Physics, Biol. Med. 2001, 13 (1), 19–27. https://doi.org/10.1007/BF02668647.

49. Borel, M.; Pastoureau, P.; Papon, J.; Madelmont, J. C.; Moins, N.; Maublant, J.; Miot-Noirault, E. Longitudinal Profiling of Articular Cartilage Degradation in Osteoarthritis by High-Resolution Magic Angle Spinning ^1^ H NMR Spectroscopy: Experimental Study in the Meniscectomized Guinea Pig Model. J. Proteome Res. 2009, 8 (5), 2594–2600. https://doi.org/10.1021/pr8009963.

50. Shet, K.; Siddiqui, S. M.; Yoshihara, H.; Kurhanewicz, J.; Ries, M.; Li, X. High-Resolution Magic Angle Spinning NMR Spectroscopy of Human Osteoarthritic Cartilage. NMR Biomed. 2012, 25 (4), 538–544. https://doi.org/10.1002/nbm.1769.

51. McIlwraith, C. W.; Frisbie, D. D.; Kawcak, C. E.; Fuller, C. J.; Hurtig, M.; Cruz, A. The OARSI Histopathology Initiative – Recommendations for Histological Assessments of Osteoarthritis in the Horse. Osteoarthr. Cartil. 2010, 18, S93–S105. https://doi.org/10.1016/J.JOCA.2010.05.031.

52. Sumner, L. W.; Amberg, A.; Barrett, D.; Beale, M. H.; Beger, R.; Daykin, C. A.; Fan, T. W.; Fiehn, O.; Goodacre, R.; Griffin, J. L.;, et al. Proposed Minimum Reporting Standards for Chemical Analysis Chemical Analysis Working Group (CAWG) Metabolomics Standards Initiative (MSI). Metabolomics 2007, 3 (3), 211–221. https://doi.org/10.1007/s11306-007-0082-2.

53. Haug, K.; Cochrane, K.; Nainala, V. C.; Williams, M.; Chang, J.; Jayaseelan, K. V.; O’Donovan, C. MetaboLights: A Resource Evolving in Response to the Needs of Its Scientific Community. Nucleic Acids Res. 2019, 48. https://doi.org/10.1093/nar/gkz1019.

54. Perez-Riverol, Y.; Csordas, A.; Bai, J.; Bernal-Llinares, M.; Hewapathirana, S.; Kundu, D. J.; Inuganti, A.; Griss, J.; Mayer, G.; Eisenacher, M.;, et al. The PRIDE Database and Related Tools and Resources in 2019: Improving Support for Quantification Data. Nucleic Acids Res. 2019, 47 (D1), D442–D450. https://doi.org/10.1093/nar/gky1106.

55. Dieterle, F.; Ross, A.; Schlotterbeck, G.; Senn, H. Probabilistic Quotient Normalization as Robust Method to Account for Dilution of Complex Biological Mixtures. Application in 1H NMR Metabonomics. 2006. https://doi.org/10.1021/AC051632C.

56. Worley, B.; Powers, R. Multivariate Analysis in Metabolomics. Curr. Metabolomics 2013, 1 (1), 92–107. https://doi.org/10.2174/2213235X11301010092.

57. Benjamini, Y.; Hochberg, Y. Controlling the False Discovery Rate: A Practical and Powerful Approach to Multiple Testing. J. R. Stat. Soc. Ser. B 1995, 57, 289–300. https://doi.org/10.2307/2346101.

58. Ogata, H.; Goto, S.; Sato, K.; Fujibuchi, W.; Bono, H.; Kanehisa, M. KEGG: Kyoto Encyclopedia of Genes and Genomes. Nucleic Acids Res. 1999, 27 (1), 29–34.

59. Szklarczyk, D.; Franceschini, A.; Wyder, S.; Forslund, K.; Heller, D.; Huerta-Cepas, J.; Simonovic, M.; Roth, A.; Santos, A.; Tsafou, K. P.;, et al. STRING V10: Protein-Protein Interaction Networks, Integrated over the Tree of Life. Nucleic Acids Res. 2015, 43 (Database issue), D447–52. https://doi.org/10.1093/nar/gku1003.

60. Salek, R. M.; Steinbeck, C.; Viant, M. R.; Goodacre, R.; Dunn, W. B. The Role of Reporting Standards for Metabolite Annotation and Identification in Metabolomic Studies. Gigascience 2013, 2 (1), 13. https://doi.org/10.1186/2047-217X-2-13.

61. Considine, E.; Salek, R. A Tool to Encourage Minimum Reporting Guideline Uptake for Data Analysis in Metabolomics. Metabolites 2019, 9 (3), 43. https://doi.org/10.3390/metabo9030043.

62. Mackay, A. R.; Ballin, M.; Pelina, M. D.; Farina, A. R.; Nason, A. M.; Hartzler, J. L.; Thorgeirsson, U. P. Effect of Phorbol Ester and Cytokines on Matrix Metalloproteinase and Tissue Inhibitor of Metalloproteinase Expression in Tumor and Normal Cell Lines. Invasion Metastasis 1992, 12 (3–4), 168–184.

63. Brama, P. A. J.; Boom, R.; DEGroot, J.; Kiers, G. H.; Weeren, P. R. Collagenase-1 (MMP-1) Activity in Equine Synovial Fluid: Influence of Age, Joint Pathology, Exercise and Repeated Arthrocentesis. Equine Vet. J. 2004, 36 (1), 34–40. https://doi.org/10.2746/0425164044864705.

64. van den Boom, R.; van der Harst, M. R.; Brommer, H.; Brama, P. A. J.; Barneveld, A.; van Weeren, P. R.; De Groot, J. Relationship between Synovial Fluid Levels of Glycosaminoglycans, Hydroxyproline and General MMP Activity and the Presence and Severity of Articular Cartilage Change on the Proximal Articular Surface of P1. Equine Vet. J. 2005, 37 (1), 19–25. https://doi.org/10.2746/0425164054406919.

65. Tseng, S.; Reddi, A. H.; Di Cesare, P. E. Cartilage Oligomeric Matrix Protein (COMP): A Biomarker of Arthritis. Biomark. Insights 2009, 4, 33–44.

66. Svala, E.; Löfgren, M.; Sihlbom, C.; Rüetschi, U.; Lindahl, A.; Ekman, S.; Skiöldebrand, E. An Inflammatory Equine Model Demonstrates Dynamic Changes of Immune Response and Cartilage Matrix Molecule Degradation in Vitro. Connect. Tissue Res. 2015, 56 (4), 315–325. https://doi.org/10.3109/03008207.2015.1027340.

67. Balakrishnan, L.; Nirujogi, R. S.; Ahmad, S.; Bhattacharjee, M.; Manda, S. S.; Renuse, S.; Kelkar, D. S.; Subbannayya, Y.; Raju, R.; Goel, R.;, et al. Proteomic Analysis of Human Osteoarthritis Synovial Fluid. Clin Proteomics 2014, 11 (1), 6. https://doi.org/10.1186/1559-0275-11-6.

68. Taylor, S. E.; Weaver, M. P.; Pitsillides, A. A.; Wheeler, B. T.; Wheeler-Jones, C. P. D.; Shaw, D. J.; Smith, R. K. W. Cartilage Oligomeric Matrix Protein and Hyaluronan Levels in Synovial Fluid from Horses with Osteoarthritis of the Tarsometatarsal Joint Compared to a Control Population. Equine Vet. J. 2006, 38 (6), 502–507.

69. Stashak, T. S.; Theoret, C. Equine Wound Management; John Wiley & Sons, 2011.

70. Peffers, M. J. Proteomic and Transcriptomic Signatures of Cartilage Ageing and Disease, University of Liverpool, 2013, Vol. PhD.

71. Lust, G.; Burton-Wurster, N.; Leipold, H. Fibronectin as a Marker for Osteoarthritis. J. Rheumatol. 1987, 14 Spec No, 28–29.

72. Murray, R. C.; Janicke, H. C.; Henson, F. M.; Goodship, A. Equine Carpal Articular Cartilage Fibronectin Distribution Associated with Training, Joint Location and Cartilage Deterioration. Equine Vet. J. 2000, 32 (1), 47–51.

73. Shikhman, A. R.; Brinson, D. C.; Valbracht, J.; Lotz, M. K. Cytokine Regulation of Facilitated Glucose Transport in Human Articular Chondrocytes. J. Immunol. 2001, 167 (12), 7001–7008. https://doi.org/10.4049/JIMMUNOL.167.12.7001.

74. Hernvann, A.; Jaffray, P.; Hilliquin, P.; Cazalet, C.; Menkes, C.-J.; Ekindjian, O. G. Interleukin-1β-Mediated Glucose Uptake by Chondrocytes. Inhibition by Cortisol. Osteoarthr. Cartil. 1996, 4 (2), 139–142. https://doi.org/10.1016/S1063-4584(05)80322-X.

75. Zhang, W.; Sun, G.; Likhodii, S.; Liu, M.; Aref-Eshghi, E.; Harper, P. E.; Martin, G.; Furey, A.; Green, R.; Randell, E.;, et al. Metabolomic Analysis of Human Plasma Reveals That Arginine Is Depleted in Knee Osteoarthritis Patients. Osteoarthr. Cartil. 2016, 24 (5), 827–834. https://doi.org/10.1016/J.JOCA.2015.12.004.

76. Hu, T.; Oksanen, K.; Zhang, W.; Randell, E.; Furey, A.; Sun, G.; Zhai, G. An Evolutionary Learning and Network Approach to Identifying Key Metabolites for Osteoarthritis. PLoS Comput. Biol. 2018, 14 (3), e1005986. https://doi.org/10.1371/journal.pcbi.1005986.

77. Abramson, S. B. Osteoarthritis and Nitric Oxide. Osteoarthr. Cartil. 2008, 16, S15– S20. https://doi.org/10.1016/S1063-4584(08)60008-4.

78. Loeser, R. F.; Carlson, C. S.; Carlo, M. Del Cole, A. Detection of Nitrotyrosine in Aging and Osteoarthritic Cartilage: Correlation of Oxidative Damage with the Presence of Interleukin-1? And with Chondrocyte Resistance to Insulin-like Growth Factor 1. Arthritis Rheum. 2002, 46 (9), 2349–2357. https://doi.org/10.1002/art.10496.

79. Lamers, R.-J. A. N.; DeGroot, J.; Spies-Faber, E. J.; Jellema, R. H.; Kraus, V. B.; Verzijl, N.; TeKoppele, J. M.; Spijksma, G. K.; Vogels, J. T. W. E.; van der Greef, J.;, et al. Identification of Disease- and Nutrient-Related Metabolic Fingerprints in Osteoarthritic Guinea Pigs. J. Nutr. 2003, 133 (6), 1776–1780. https://doi.org/10.1093/jn/133.6.1776.

80. Zhai, G.; Wang-Sattler, R.; Hart, D. J.; Arden, N. K.; Hakim, A. J.; Illig, T.; Spector, T. D. Serum Branched-Chain Amino Acid to Histidine Ratio: A Novel Metabolomic Biomarker of Knee Osteoarthritis. Ann. Rheum. Dis. 2010, 69 (6), 1227–1231. https://doi.org/10.1136/ard.2009.120857.

81. Phang, J. M.; Liu, W.; Hancock, C. N.; Fischer, J. W. Proline Metabolism and Cancer: Emerging Links to Glutamine and Collagen. Curr. Opin. Clin. Nutr. Metab. Care 2015, 18 (1), 71–77. https://doi.org/10.1097/MCO.0000000000000121.

82. Milner, J. M.; Elliott, S.-F.; Cawston, T. E. Activation of Procollagenases Is a Key Control Point in Cartilage Collagen Degradation: Interaction of Serine and Metalloproteinase Pathways. Arthritis Rheum. 2001, 44 (9), 2084–2096. https://doi.org/10.1002/1529-0131(200109)44:9<2084::AID-ART359>3.0.CO;2-R.

83. Milner, J. M.; Rowan, A. D.; Cawston, T. E.; Young, D. A. Metalloproteinase and Inhibitor Expression Profiling of Resorbing Cartilage Reveals Pro-Collagenase Activation as a Critical Step for Collagenolysis. Arthritis Res. Ther. 2006, 8 (5), R142. https://doi.org/10.1186/ar2034.

84. Gupta, S.; Biswas, A.; Akhter, M. S.; Krettler, C.; Reinhart, C.; Dodt, J.; Reuter, A.; Philippou, H.; Ivaskevicius, V.; Oldenburg, J. Revisiting the Mechanism of Coagulation Factor XIII Activation and Regulation from a Structure/Functional Perspective. Sci. Rep. 2016, 6 (1), 30105. https://doi.org/10.1038/srep30105.

85. Johnson, K.; Hashimoto, S.; Lotz, M.; Pritzker, K.; Terkeltaub, R. Interleukin-1 Induces pro-Mineralizing Activity of Cartilage Tissue Transglutaminase and Factor XIIIa. Am. J. Pathol. 2001, 159 (1), 149–163. https://doi.org/10.1016/S0002-9440(10)61682-3.

86. Sanchez, C.; Deberg, M. A.; Bellahcène, A.; Castronovo, V.; Msika, P.; Delcour, J. P.; Crielaard, J. M.; Henrotin, Y. E. Phenotypic Characterization of Osteoblasts from the Sclerotic Zones of Osteoarthritic Subchondral Bone. Arthritis Rheum. 2008, 58 (2), 442–455. https://doi.org/10.1002/art.23159.

87. Day, J. S.; Van Der Linden, J. C.; Bank, R. A.; Ding, M.; Hvid, I.; Sumner, D. R.; Weinans, H. Adaptation of Subchondral Bone in Osteoarthritis. Biorheology 2004, 41 (3–4), 359–368.

88. Nurminskaya, M.; Magee, C.; Nurminsky, D.; Linsenmayer, T. F. Plasma Transglutaminase in Hypertrophic Chondrocytes: Expression and Cell-Specific Intracellular Activation Produce Cell Death and Externalization. J. Cell Biol. 1998, 142 (4), 1135–1144.

89. Rosenthal, A. K.; Masuda, I.; Gohr, C. M.; Derfus, B. A.; Le, M. The Transglutaminase, Factor XIIIA, Is Present in Articular Chondrocytes. Osteoarthr. Cartil. 2001, 9 (6), 578–581. https://doi.org/10.1053/JOCA.2000.0423.

90. Lee, J. Y.; Kang, M. J.; Choi, J. Y.; Park, J. S.; Park, J. K.; Lee, E. Y.; Lee, E. B.; Pap, T.; Yi, E. C.; Song, Y. W. Apolipoprotein B Binds to Enolase-1 and Aggravates Inflammation in Rheumatoid Arthritis. Ann. Rheum. Dis. 2018, annrheumdis-2018-213444. https://doi.org/10.1136/annrheumdis-2018-213444.

91. Sánchez-Enríquez, S.; Torres-Carrillo, N. M.; Vázquez-Del Mercado, M.; Salgado-Goytia, L.; Rangel-Villalobos, H.; Muñoz-Valle, J. F. Increase Levels of Apo-A1 and Apo B Are Associated in Knee Osteoarthritis: Lack of Association with VEGF-460 T/C and +405 C/G Polymorphisms. Rheumatol. Int. 2008, 29 (1), 63–68. https://doi.org/10.1007/s00296-008-0633-5.

92. Roughley, P. J.; White, R. J.; Cs-Szabó, G.; Mort, J. S. Changes with Age in the Structure of Fibromodulin in Human Articular Cartilage. Osteoarthr. Cartil. 1996, 4 (3), 153–161. https://doi.org/10.1016/S1063-4584(96)80011-2.

93. Wadhwa, S.; Embree, M. C.; Kilts, T.; Young, M. F.; Ameye, L. G. Accelerated Osteoarthritis in the Temporomandibular Joint of Biglycan/Fibromodulin Double-Deficient Mice. Osteoarthr. Cartil. 2005, 13 (9), 817–827. https://doi.org/10.1016/J.JOCA.2005.04.016.

94. Ameye, L.; Aria, D.; Jepsen, K.; Oldberg, A.; Xu, T.; Young, M. F. Abnormal Collagen Fibrils in Tendons of Biglycan/Fibromodulin-Deficient Mice Lead to Gait Impairment, Ectopic Ossification, and Osteoarthritis. FASEB J. 2002, 16 (7), 673– 680. https://doi.org/10.1096/fj.01-0848com.

95. Swift, J.; Discher, D. E. The Nuclear Lamina Is Mechano-Responsive to ECM Elasticity in Mature Tissue. J. Cell Sci. 2014, 127 (Pt 14), 3005–3015. https://doi.org/10.1242/jcs.149203.

96. Attur, M.; Ben-Artzi, A.; Yang, Q.; Al-Mussawir, H. E.; Worman, H. J.; Palmer, G.; Abramson, S. B. Perturbation of Nuclear Lamin A Causes Cell Death in Chondrocytes. Arthritis Rheum. 2012, 64 (6), 1940–1949. https://doi.org/10.1002/art.34360.

97. de Figueroa, P. L.; Nogueira-Recalde, U.; Osorio, F.; Lotz, M.; Lopez-Otin, C.; Blanco, F. J.; Carames, B. Deficient Autophagy Induces Lamin a/C Accumulation in Aging and Osteoarthritis. Am. Coll. Rheumatol. 2017, 69 (Supplement 10).

98. Ivaska, J.; Pallari, H.-M.; Nevo, J.; Eriksson, J. E. Novel Functions of Vimentin in Cell Adhesion, Migration, and Signaling. Exp. Cell Res. 2007, 313 (10), 2050–2062. https://doi.org/10.1016/J.YEXCR.2007.03.040.

99. Langelier, E.; Suetterlin, R.; Hoemann, C. D.; Aebi, U.; Buschmann, M. D. The Chondrocyte Cytoskeleton in Mature Articular Cartilage: Structure and Distribution of Actin, Tubulin, and Vimentin Filaments. J. Histochem. Cytochem. 2000, 48 (10), 1307–1320. https://doi.org/10.1177/002215540004801002.

100. Lambrecht, S.; Verbruggen, G.; Verdonk, P. C. M.; Elewaut, D.; Deforce, D. Differential Proteome Analysis of Normal and Osteoarthritic Chondrocytes Reveals Distortion of Vimentin Network in Osteoarthritis. Osteoarthr. Cartil. 2008, 16 (2), 163–173. https://doi.org/10.1016/J.JOCA.2007.06.005.

101. Lotz, M.; Martel-Pelletier, J.; Christiansen, C.; Brandi, M.-L.; Bruyère, O.; Chapurlat, R.; Collette, J.; Cooper, C.; Giacovelli, G.; Kanis, J. A.;, et al. Value of Biomarkers in Osteoarthritis: Current Status and Perspectives. Ann. Rheum. Dis. 2013, 72 (11), 1756–1763. https://doi.org/10.1136/annrheumdis-2013-203726.

102. Marshall, D. D.; Powers, R. Beyond the Paradigm: Combining Mass Spectrometry and Nuclear Magnetic Resonance for Metabolomics. Prog. Nucl. Magn. Reson. Spectrosc. 2017, 100, 1–16. https://doi.org/10.1016/J.PNMRS.2017.01.001.

103. Parker, C. E.; Borchers, C. H. Mass Spectrometry Based Biomarker Discovery, Verification, and Validation - Quality Assurance and Control of Protein Biomarker Assays. Mol. Oncol. 2014, 8 (4), 840–858. https://doi.org/10.1016/j.molonc.2014.03.006.

104. Caterson, B.; Baker, J. R.; Christnerg, J. E.; Leell, Y.; Lentzn, M. Monoclonal Antibodies as Probes for Determining the Microheterogeneity of the Link Proteins of Cartilage Proteoglycan. J. Biol. Chem. 1985, 260 (19).

