## Supplementary Information for "*Ex-Vivo* Equine Cartilage Explant Osteoarthritis Model - A Metabolomics and Proteomics Study"

*corresponding author

**Corresponding author email address**

### Contents

**Liquid Chromatography Tandem Mass Spectrometry - Detailed Methods**

**Figure S1.** Five post mortem equine metacarpophalangeal joints used for *ex-vivo* cartilage culture.

**Figure S2.** Experimental design for *ex-vivo* equine cartilage culture +/- TNF-α/IL-1β treatment. MCP = metacarpophalangeal.

**Figure S3.** 1D $^1$H nuclear magnetic resonance spectral quantile plots of cartilage - 8 days in control media, cartilage - 8 days in TNF-α/IL-1β treated media, control media (all time points combined) and TNF-α/IL-1β treated media (all time points combined).

**Figure S4.** Representative culture media ion chromatograms of combined time points for control and TNF-α/IL-1β treated equine *ex-vivo* cartilage explants using a 60 min liquid chromatography gradient.

**Figure S5.** Number of proteins identified within culture media for (A) combined and (B) individual sample time points for controls and TNF-α/IL-1β treated *ex-vivo* equine cartilage. Combined

**Figure S6.** PC1 RMS (Principal component 1 root mean square) values for the 25 components with the highest magnitude for differentially abundant proteins present within culture media at (A) 0-2 days, (B) 3-5 days and (C) 6-8 days following TNF-α/IL-1β treatment of *ex-vivo* equine cartilage. n=5 for each time point. RMS: High = high in treatment with respect to control, Low = low in treatment with respect to control.

**Figure S7.** Full Protein Gel Image used in Figure 7. Silver stain identifying media protein profiles (combined for all time points) following incubation of *ex-vivo* equine cartilage for control and TNF-α/IL-1β treated samples.

**Figure S8.** Principal component analyses of semi-tryptic peptide profiles within culture media of control and TNF-α/IL-1β treated *ex-vivo* equine cartilage at 0-2 days, 3-5 days and 6-8 days.

**Table S1.** Proteins identified within culture media of control and TNF-α/IL-1β treated *ex-vivo* equine cartilage time points.

**Liquid Chromatography Tandem Mass Spectrometry - Detailed Methods**

Tryptic digests were diluted 5-fold in 0.1% (v/v) trifluoroacetic acid (TFA) and 3% (v/v) acetonitrile and analysed individually, in a random order, via liquid chromatography tandem mass spectrometry (LC-MS/MS) using a 60 min liquid chromatography (LC) gradient. A Q Exactive™ HF quadrupole-Orbitrap mass spectrometer (Thermo Scientific, Hemel Hempstead, UK) coupled to a Dionex Ultimate 3000 RSLC nano-liquid chromatograph (Thermo Scientific) was used for data-dependent LC-MS/MS analyses. Digests were loaded onto a trapping column (Acclaim PepMap 100, C18, 20 mm x 75 µm) using a loading buffer of 0.1% (v/v) TFA and 2% (v/v) acetonitrile in water for 3 min at a flow rate of 5 µl $min^{-1}$. The trapping column was then set in-line with an analytical column (Easy-Spray PepMap® C18, 15 cm x 75 µm, 2 µm) with peptide elution carried out using a linear gradient of 96.2% A (0.1% (v/v) formic acid):3.8% B (0.1 % (v/v) formic acid in water:acetonitrile (80:20) (v/v)) to 50% A:50% B over 30 min at a flow rate of 300 nl $min^{-1}$, followed by washing at 1 % A:99% B for 5 min and re-equilibration of the column to starting conditions. Sample digests were interspersed with 30 min blanks (97% (v/v) high performance liquid chromatography grade $H_20$ (VWR International), 2.9% acetonitrile and 0.1% TFA. The Q Exactive™ was operated in data dependent positive (ESI+) mode with survey scans between *m/z* 300-2000 acquired at a mass resolution of 70,000 (full width at half maximum) at *m/z* 200 after accumulation of ions to $1x10^6$ target value based on predictive automatic gain control values from the previous full scan. The 10 most intense precursor ions with charge states of between 2+ and 5+ were selected for MS/MS with an isolation window of 2 *m/z* units. Higher-energy collisional dissociation (HCD) was used to fragment peptides using normalised collision energy of 30% with a maximum injection time of 100 ms. Dynamic exclusion of *m/z* values to prevent repeated fragmentation of the same peptide was used with an exclusion time of 20 s.

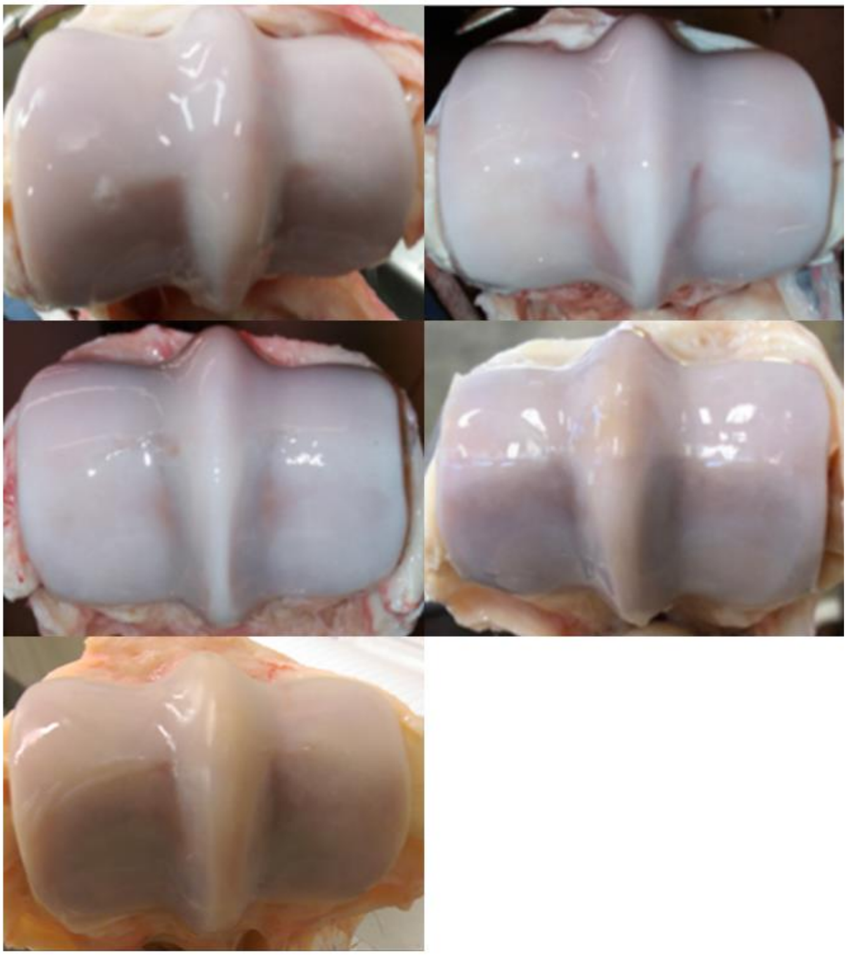

**Figure S1.** Five post mortem equine metacarpophalangeal joints used for *ex-vivo* cartilage culture. Cartilage collected from all joints was considered macroscopically normal with a score of 0 according to the OARSI histopathology initiative scoring system for horses.

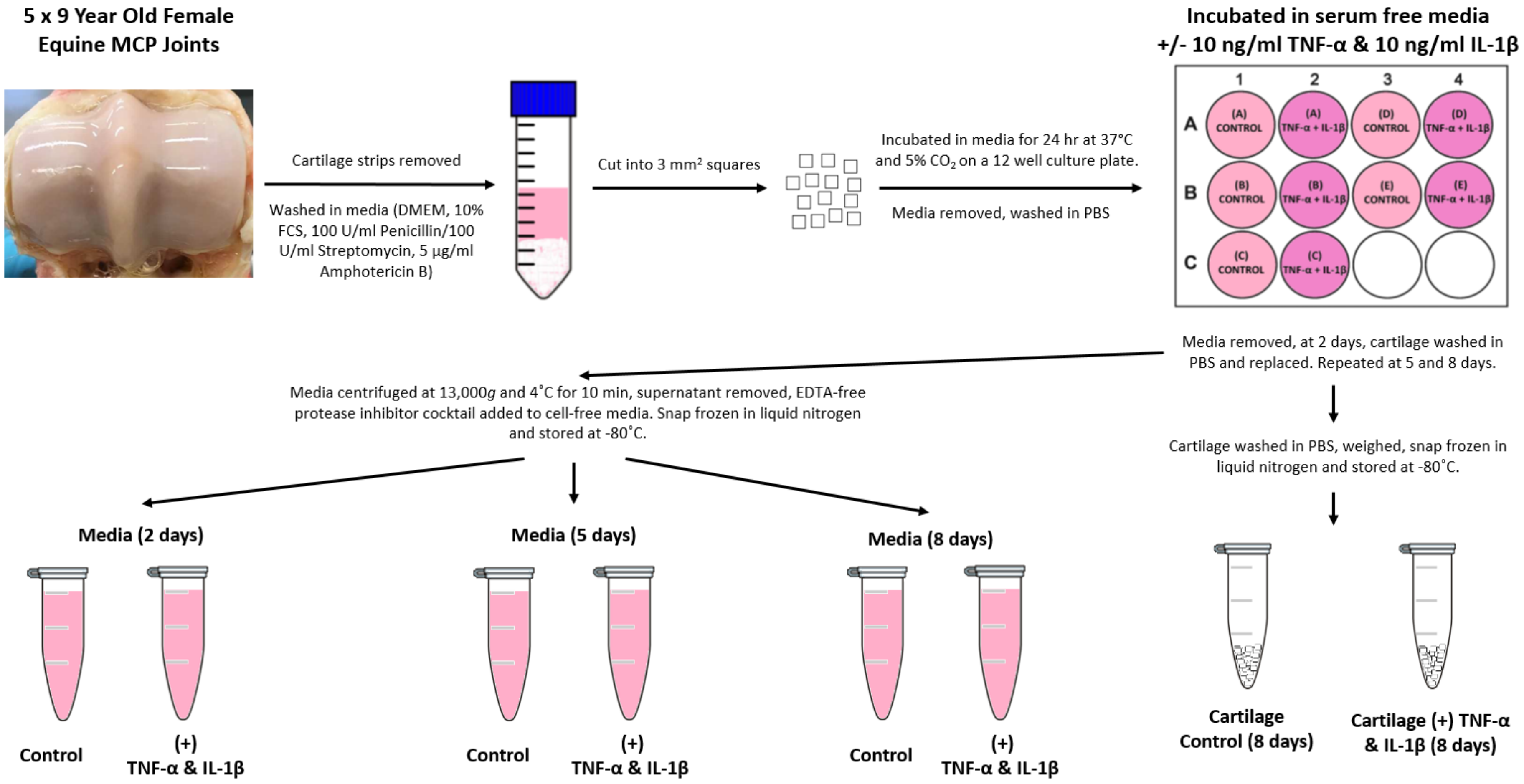


**Figure S2.** Experimental design for *ex-vivo* equine cartilage culture +/- TNF-α/IL-1β treatment. MCP = metacarpophalangeal.

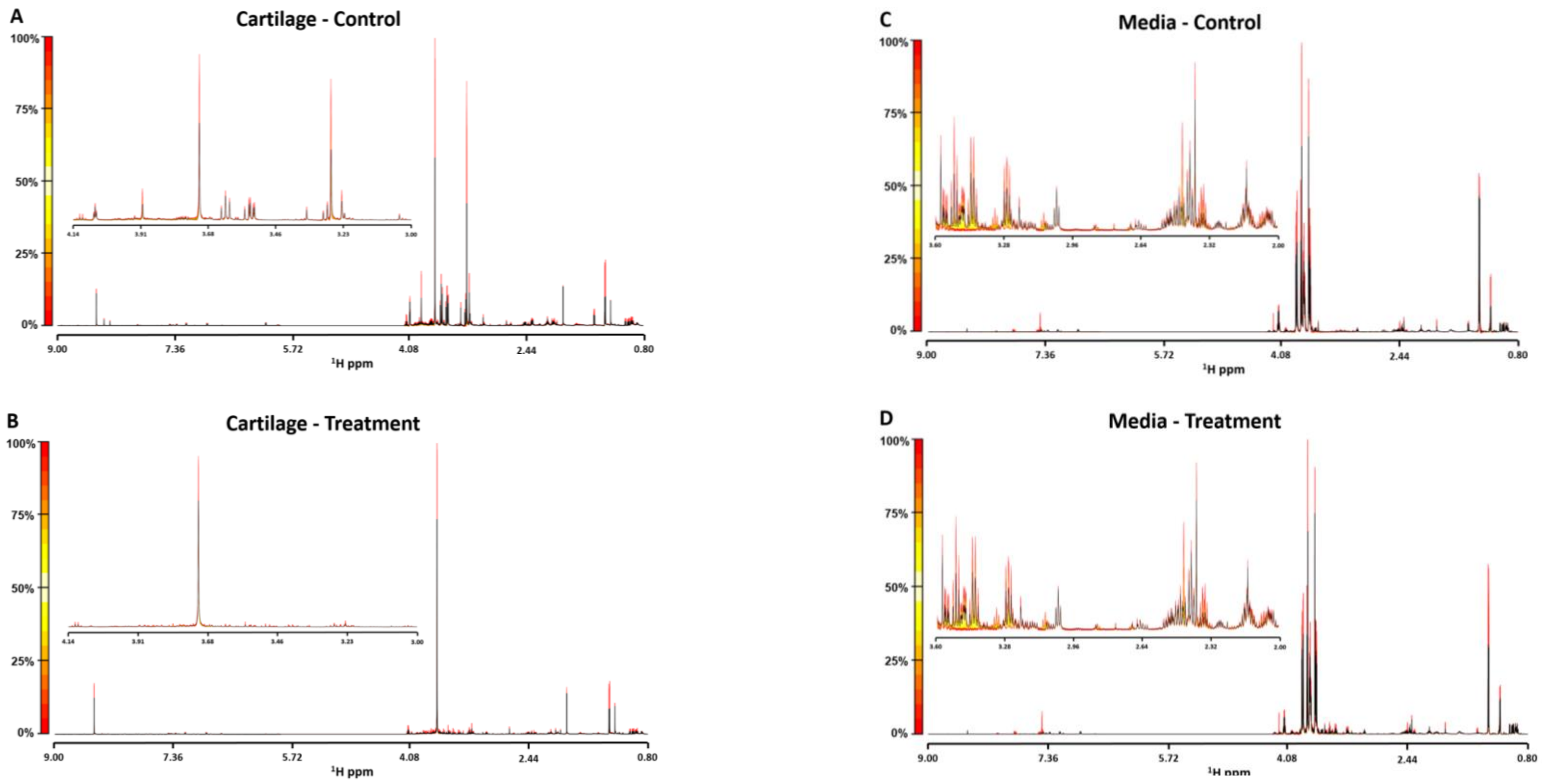


**Figure S3.** 1D $^1$H nuclear magnetic resonance spectral quantile plots of (A) cartilage, 8 days in control media (B) cartilage, 8 days in TNF-α/IL-1β treated media (C) control media (all time points combined) and (D) TNF-α/IL-1β treated media (all time points combined). The median spectral plot is depicted by a black line and variation from the median shown as a yellow to red scale for both the full spectral range (9.00-0.80 ppm) and more detailed regions (4.14-3.00 ppm for cartilage and 3.60-2.00 ppm for media). Control (n=5) and TNF-α/IL-1β treatment (n=5).

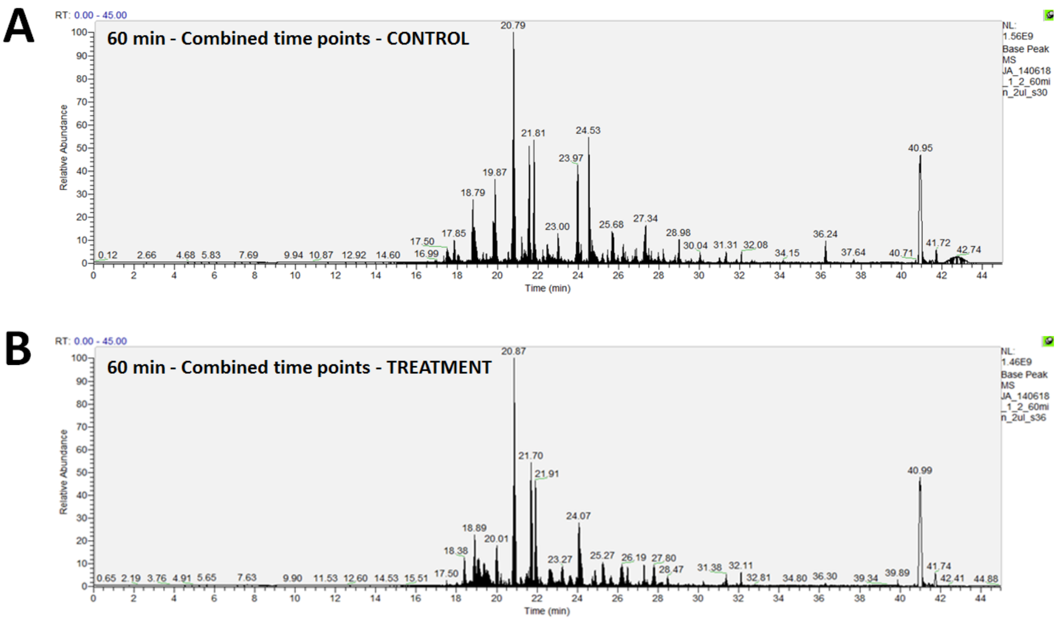


**Figure S4.** Representative culture media ion chromatograms of combined time points for (A) control and (B) TNF-α/IL-1β treated equine *ex-vivo* cartilage explants using a 60 min liquid chromatography gradient.

A

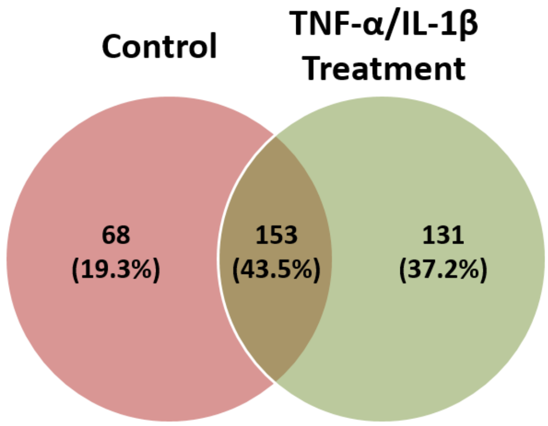


B

| | No. Proteins Identified/Media Sample | |
|---|---|---|
| | Mean | Median |
| Control | 124 | 112 |
| TNF-α/IL-1β Treatment | 189 | 191 |

**Figure S5.** Number of proteins identified within culture media for (A) combined and (B) individual sample time points for controls and TNF-α/IL-1β treated *ex-vivo* equine cartilage. Combined time points, n=5/group, Individual time points, n=15/group.

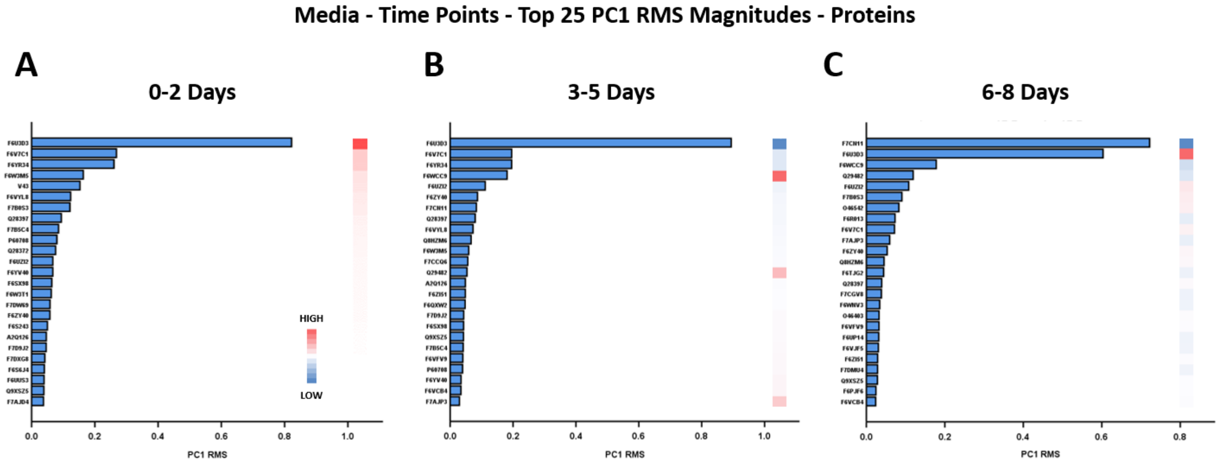


**Figure S6.** PC1 RMS (Principal component 1 root mean square) values for the 25 components with the highest magnitude for differentially abundant proteins present within culture media at (A) 0-2 days, (B) 3-5 days and (C) 6-8 days following TNF-α/IL-1β treatment of *ex-vivo* equine cartilage. n=5 for each time point. RMS: High = high in treatment with respect to control, Low = low in treatment with respect to control.

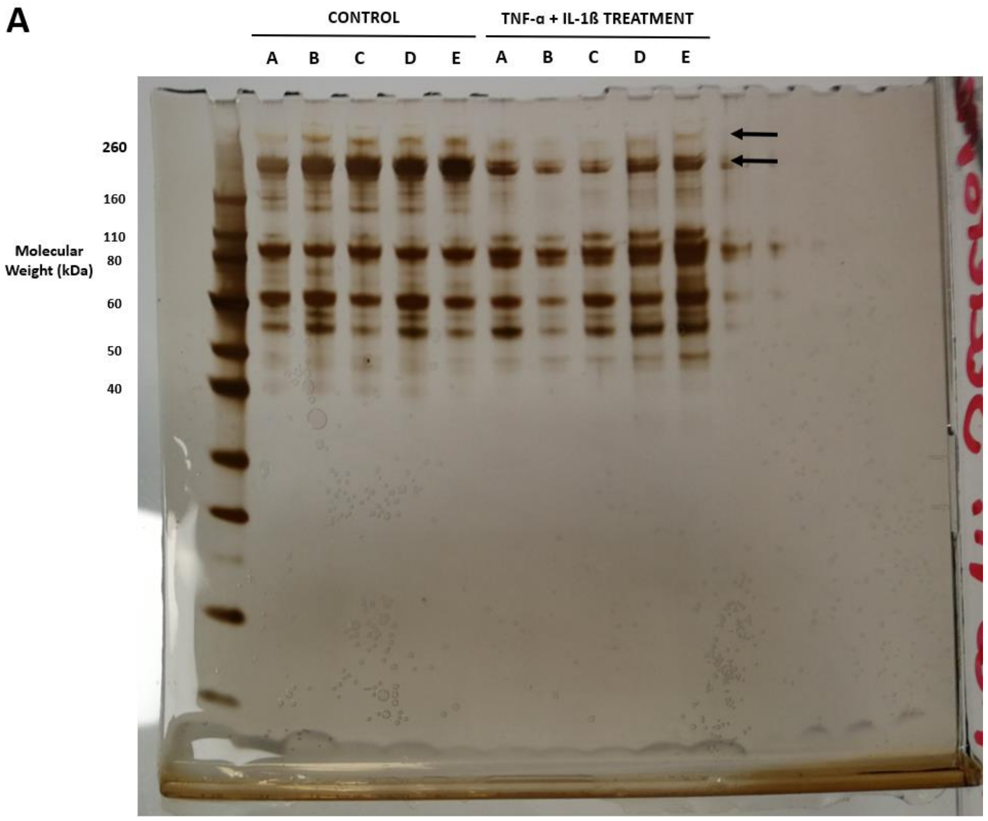


**Figure S7.** Full Protein Gel Image used in Figure 7. Silver stain identifying media protein profiles (combined for all time points) following incubation of *ex-vivo* equine cartilage for control and TNF-α/IL-1β treated samples. Arrows indicate differentially secreted proteins at approximately 160-260 kDa and 260 kDa.

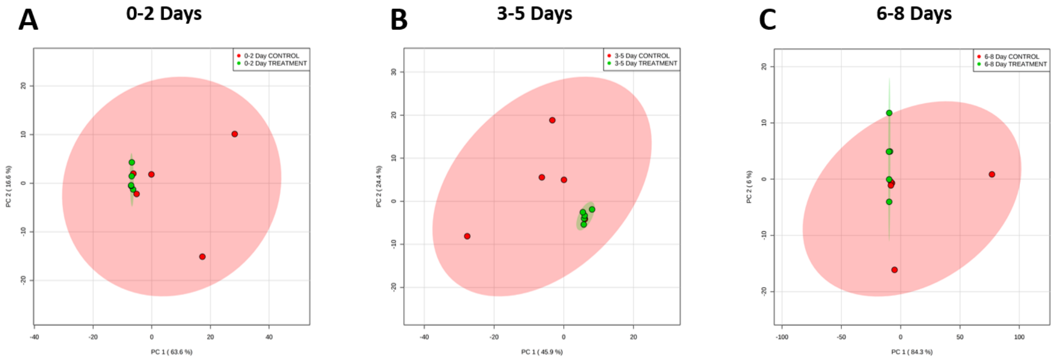


**Figure S8.** Principal component analyses of semi-tryptic peptide profiles within culture media of control (red, n=5) and TNF-α/IL-1β treated (green, n=5) *ex-vivo* equine cartilage at (A) 0-2 days, (B) 3-5 days and (C) 6-8 days.

**Table S1.** Proteins identified within culture media of control and TNF-α/IL-1β treated *ex-vivo* equine cartilage.

| Media Samples Identified In | Accession Number | Family/Subfamily |
|---|---|---|
| Control Only | F6PLF6 | 2-Phosphoxylose Phosphatase 1 |
| Control Only | F6R1X9 | Actin, Alpha Cardiac Muscle 1 |
| Control Only | F7CZ92 | Actin, Alpha Skeletal Muscle |
| Control Only | F7CW82 | Actin, Aortic Smooth Muscle |
| Control Only | F6Z8K0 | Actin, Gamma-Enteric Smooth Muscle |
| Control Only | F7C2Y5 | Adipocyte Enhancer-Binding Protein 1 |
| Control Only | F6YRA8 | ATP-Dependent rna Helicase Ddx60-Related |
| Control Only | F7CPZ3 | Calsyntenin 1 |
| Control Only | F6YQD9 | Carboxypeptidase E |
| Control Only | F7CCP9 | Cartilage Intermediate Layer Protein 2 |
| Control Only | F6VF11 | Chordin-Like Protein 2 |
| Control Only | F6VP03 | Collagen Type IX Alpha 3 Chain |
| Control Only | F6VBF7 | Collagen Type XI Alpha 1 Chain |
| Control Only | F6U661 | Collagen Type XI Alpha 1 Chain |
| Control Only | F6VAW6 | Collagen Type XI Alpha 1 Chain |
| Control Only | F7BQD6 | Complement C1S Subcomponent |
| Control Only | F6ZR63 | Complement Factor H |
| Control Only | F7AU56 | Complement Factor I |
| Control Only | F6Z7T9 | Ectonucleotide Pyrophosphatase/Phosphodiesterase 2 |
| Control Only | F6Z7X8 | Ectonucleotide Pyrophosphatase/Phosphodiesterase Family Member 2 |

| | | |
|---|---|---|
| Control Only | F6Q4R1 | Elongation Factor 1-Alpha 2 |
| Control Only | F6WD07 | Fibulin 7 |
| Control Only | F7DHZ1 | Gelsolin |
| Control Only | F7E2D1 | Gelsolin |
| Control Only | F7D7T6 | Heat Shock 70 KDa Protein 1-Like |
| Control Only | F6TWM4 | Histone H2B |
| Control Only | F6WFI8 | Histone H2B Type 1-A |
| Control Only | F7ASE9 | Histone H2B Type 1-C/E/F/G/I |
| Control Only | F6VUX0 | Histone H2B Type 1-D |
| Control Only | F7CJ51 | Histone H2B Type 1-D |
| Control Only | F7E1X9 | Histone H2B Type 1-H |
| Control Only | F6VYH8 | Histone H2B Type 1-M |
| Control Only | F7DLL0 | Histone H2B Type 1-N |
| Control Only | F6PWV1 | Histone H2B Type 2-F |
| Control Only | Q95LJ1 | Insulin-Like Growth Factor-Binding Protein 7 |
| Control Only | F6SSR0 | Integrin Beta-Like Protein 1 |
| Control Only | F6QBB1 | Inter-Alpha-Trypsin Inhibitor Heavy Chain H5 |
| Control Only | F6ZEQ3 | Keratin, Type I Cytoskeletal 16 |
| Control Only | F7AGY4 | Keratin, Type Ii Cytoskeletal 6B-Related |
| Control Only | F6ST15 | Keratin-87 Protein-Related |
| Control Only | F6WZ08 | Laminin Subunit Gamma-2 |
| Control Only | Q8HZI9 | Laminin Subunit Gamma-2 |
| Control Only | F6W6E8 | Latent Transforming Growth Factor Beta Binding Protein 1 |
| Control Only | F6W6Y4 | Latent-Transforming Growth Factor Beta-Binding Protein 1 |
| Control Only | F6Z0F8 | Latent-Transforming Growth Factor Beta-Binding Protein 3 |
| Control Only | F6UXM7 | Lysyl Oxidase Homolog 3 |
| Control Only | O02722 | Metalloproteinase Inhibitor 1 |
| Control Only | F7BEE7 | Metalloproteinase Inhibitor 2 |
| Control Only | F7AEL9 | Neuron Derived Neurotrophic Factor |
| Control Only | F7BQX4 | Peptidylglycine Alpha-Amidating Monooxygenase |
| Control Only | F7BR00 | Peptidylglycine Alpha-Amidating Monooxygenase |
| Control Only | F6ZCC8 | Peptidyl-Glycine Alpha-Amidating Monooxygenase |
| Control Only | F7D1R1 | Phosphoglycerate Kinase 1 |
| Control Only | F6VTS1 | Phospholipase A1 Member A |
| Control Only | F7CR03 | Phospholipid Transfer Protein |
| Control Only | F7BKE1 | Pigment Epithelium-Derived Factor |
| Control Only | F6R5Y2 | Plasma Serine Protease Inhibitor |
| Control Only | F6V2X7 | Procollagen-Lysine,2-Oxoglutarate 5-Dioxygenase 1 |
| Control Only | F6VTD3 | Proteoglycan 4 |
| Control Only | F6XL03 | Scrapie-Responsive Protein 1 |
| Control Only | F6XVP2 | Sparc-Related Modular Calcium-Binding Protein 1 |

| | | |
|---|---|---|
| Control Only | F6WQ44 | Sulfhydryl Oxidase |
| Control Only | F6WR95 | Sulfhydryl Oxidase 1 |
| Control Only | O19011 | Transforming Growth Factor Beta-1 |
| Control Only | F7APU2 | Uncharacterized Protein |
| Control Only | F6XLT6 | Vascular Endothelial Growth Factor A |
| Control Only | Q9GKR0 | Vascular Endothelial Growth Factor A |
| Control Only | F7AZP8 | Xylosyltransferase 1 |
| Treatment Only | F6T0A6 | 10 KDa Heat Shock Protein, Mitochondrial |
| Treatment Only | F6SP02 | 14-3-3 Protein Theta |
| Treatment Only | F7BFM4 | 60S Ribosomal Protein L10A |
| Treatment Only | F6T1A4 | 60S Ribosomal Protein L12 |
| Treatment Only | F6Z421 | 60S Ribosomal Protein L12 |
| Treatment Only | F7D917 | 6-Phosphogluconate Dehydrogenase, Decarboxylating |
| Treatment Only | F6YG82 | 78 KDa Glucose-Regulated Protein |
| Treatment Only | F6PH57 | Actin-Related Protein 2/3 Complex Subunit 4 |
| Treatment Only | F6W354 | Actin-Related Protein 3 |
| Treatment Only | F6UFV3 | Adenosine Kinase |
| Treatment Only | F6TL52 | Adenylate Kinase Isoenzyme 1 |
| Treatment Only | F6SRP7 | Adenylyl Cyclase-Associated Protein 1 |
| Treatment Only | F7AJD4 | AHNAK Nucleoprotein |
| Treatment Only | F7CBN0 | Alcohol Dehydrogenase (NADP+)-Like Protein |
| Treatment Only | P19854 | Alcohol Dehydrogenase Class-3 |
| Treatment Only | F7DMQ3 | Aldose Reductase |
| Treatment Only | F7E419 | Annexin A8 |
| Treatment Only | F6ZB53 | Apurinic Or Apyrimidinic Site) Lyase;Apex1;Ortholog |
| Treatment Only | F6ZXT0 | Atpase Inhibitor, Mitochondrial |
| Treatment Only | F7C555 | Bifunctional Purine Biosynthesis Protein Purh |
| Treatment Only | F6YIU8 | C-1-Tetrahydrofolate Synthase, Cytoplasmic |
| Treatment Only | F7D854 | Catalase |
| Treatment Only | F6YF95 | Cellular Nucleic Acid-Binding Protein |
| Treatment Only | F7CR22 | Coatomer Subunit Delta |
| Treatment Only | F7DXG8 | Cofilin-1 |
| Treatment Only | F6ZNX3 | Collagen Alpha-1 |
| Treatment Only | F7DEW5 | Coronin-1C |
| Treatment Only | F6S9Z1 | C-X-C Motif Chemokine 5 |
| Treatment Only | Q8MIN2 | C-X-C Motif Chemokine 5 |
| Treatment Only | F6XJR7 | Cysteine And Glycine-Rich Protein 1 |
| Treatment Only | F7B5P1 | Cytosolic Non-Specific Dipeptidase |
| Treatment Only | F7B320 | Dihydropyrimidinase-Related Protein 2 |
| Treatment Only | A2Q127 | Elongation Factor 1-Gamma |
| Treatment Only | F6XTN1 | Eukaryotic Initiation Factor 4A-I |
| Treatment Only | F6YYS2 | Eukaryotic Initiation Factor 4A-Ii |
| Treatment Only | F7BPT4 | Ezrin |

| | | |
|---|---|---|
| Treatment Only | F6ZYU6 | Far Upstream Element-Binding Protein 1 |
| Treatment Only | F7A984 | Far Upstream Element-Binding Protein 2 |
| Treatment Only | F7CS60 | Fibronectin |
| Treatment Only | F7D2D9 | Flavin Reductase |
| Treatment Only | F6RKC6 | Four and A Half Lim Domains Protein 1 |
| Treatment Only | F7CL92 | Glutamine--Fructose-6-Phosphate Aminotransferase [Isomerizing] 1 |
| Treatment Only | F6VSN2 | Glutathione S-Transferase P |
| Treatment Only | F7BU09 | Glutathione S-Transferase Pi 3 |
| Treatment Only | F7DZD2 | Glutathione S-Transferase Theta-1 |
| Treatment Only | F6RV94 | Glycogen Phosphorylase, Liver Form |
| Treatment Only | B7UBT6 | Growth-Regulated Alpha Protein |
| Treatment Only | F6R8V8 | Growth-Regulated Alpha Protein |
| Treatment Only | A2Q0Z1 | Heat Shock Cognate 71 Kda Protein |
| Treatment Only | F7AUW2 | Heat Shock Cognate 71 Kda Protein |
| Treatment Only | F7E3Y7 | Heat Shock Protein Beta-1 |
| Treatment Only | F6TF34 | Heme Oxygenase 1 |
| Treatment Only | F7A868 | Heterogeneous Nuclear Ribonucleoprotein A1 |
| Treatment Only | F6YZ44 | Heterogeneous Nuclear Ribonucleoprotein A1-Related |
| Treatment Only | F6Z0C6 | Heterogeneous Nuclear Ribonucleoprotein D |
| Treatment Only | F6WHS3 | Heterogeneous Nuclear Ribonucleoprotein D0 |
| Treatment Only | F6UQU2 | Heterogeneous Nuclear Ribonucleoprotein D-Like |
| Treatment Only | F6Q4Q1 | Heterogeneous Nuclear Ribonucleoprotein K |
| Treatment Only | F6RTD6 | Heterogeneous Nuclear Ribonucleoprotein L |
| Treatment Only | F7BA08 | Heterogeneous Nuclear Ribonucleoprotein R |
| Treatment Only | F6VYB1 | Heterogeneous Nuclear Ribonucleoproteins A2/B1 |
| Treatment Only | F6S2F8 | Histone H1.2 |
| Treatment Only | F7BCH1 | Inhibin Beta A Chain |
| Treatment Only | P55102 | Inhibin Beta A Chain |
| Treatment Only | F6QJ27 | Isochorismatase Domain-Containing Protein 1 |
| Treatment Only | F6R7J2 | Isocitrate Dehydrogenase [Nadp] Cytoplasmic |
| Treatment Only | F6ZTT5 | Keratin 33B |
| Treatment Only | F6PJX6 | Keratin, Type I Cytoskeletal 13 |
| Treatment Only | F6VNP6 | Keratin, Type II Cuticular Hb5 |
| Treatment Only | F6SHJ8 | Keratin, Type II Cytoskeletal 2 Epidermal |
| Treatment Only | F7C7Y1 | Keratin, Type II Cytoskeletal 73 |
| Treatment Only | F7B232 | Lambda-Crystallin Homolog |
| Treatment Only | F7DQB9 | Lim and Cysteine-Rich Domains Protein 1 |
| Treatment Only | F6PSB8 | Lim and Sh3 Domain Protein 1 |
| Treatment Only | C6L1J5 | L-Lactate Dehydrogenase B Chain |
| Treatment Only | F7D144 | Macrophage-Capping Protein |
| Treatment Only | F6SAI2 | Matrilin-4 |
| Treatment Only | F6WTM6 | Matrix Metalloproteinase 1 |

| | | |
|---|---|---|
| Treatment Only | Q9XSZ5 | Matrix Metalloproteinase 1 |
| Treatment Only | F6U999 | Mesencephalic Astrocyte-Derived Neurotrophic Factor |
| Treatment Only | Q9TUL9 | Metalloproteinase Inhibitor 3 |
| Treatment Only | F7DM22 | Metallothionein |
| Treatment Only | P02801 | Metallothionein-1B |
| Treatment Only | F6VKN2 | Metallothionein-2 |
| Treatment Only | F6PVJ6 | Mimecan |
| Treatment Only | F7D7Z9 | Multifunctional Protein Ade2 |
| Treatment Only | F6X3K0 | Nucleolin |
| Treatment Only | F6X6A6 | Peptidyl-Prolyl Cis-Trans Isomerase-Related |
| Treatment Only | F7AXI9 | Peroxiredoxin-6 |
| Treatment Only | F7DQS6 | Phosphoglycerate Mutase 1 |
| Treatment Only | F7AGA5 | Plastin-3 |
| Treatment Only | F6WPZ3 | Plectin |
| Treatment Only | F6WQ73 | Plectin |
| Treatment Only | F6WQF0 | Plectin |
| Treatment Only | F6WQW5 | Plectin |
| Treatment Only | F6WXR4 | Plectin |
| Treatment Only | F6WRA4 | Plectin |
| Treatment Only | F6TQ92 | Polyadenylate-Binding Protein 1 |
| Treatment Only | F6UGL6 | Polypyrimidine Tract-Binding Protein 1 |
| Treatment Only | F6TEI8 | Profilin-1 |
| Treatment Only | F6UJ33 | Profilin-1 |
| Treatment Only | F6VS95 | Protein Disulfide-Isomerase A6 |
| Treatment Only | F6WPB4 | Protein Dj-1 |
| Treatment Only | F6UNB9 | Protein S100-A1 |
| Treatment Only | F7BND9 | Protein S100-A11 |
| Treatment Only | F7CB04 | Protein Transport Protein Sec23A |
| Treatment Only | F7C108 | Protein-Lysine 6-Oxidase |
| Treatment Only | F6WB51 | Quinone Oxidoreductase |
| Treatment Only | F6Z4J4 | Rab Gdp Dissociation Inhibitor Beta |
| Treatment Only | F6YRC5 | Ras Gtpase-Activating-Like Protein Iqgap1 |
| Treatment Only | F6XJB9 | RNA-Binding Motif Protein, X Chromosome |
| Treatment Only | F7ASU6 | Selenium Binding Protein 1 |
| Treatment Only | F7CI32 | Selenium Binding Protein 1 |
| Treatment Only | F7DKR3 | Selenium-Binding Protein 1 |
| Treatment Only | F6XS05 | S-Formylglutathione Hydrolase |
| Treatment Only | F6PZ47 | Staphylococcal Nuclease Domain-Containing Protein 1 |
| Treatment Only | F7E1X2 | Stress-Induced-Phosphoprotein 1 |
| Treatment Only | F6XC16 | Synaptic Vesicle Membrane Protein Vat-1 Homolog |
| Treatment Only | F6UMQ4 | Transforming Growth Factor Beta Induced |
| Treatment Only | F6QXN5 | Transgelin-2 |
| Treatment Only | F6TZS9 | Triosephosphate Isomerase |

| | | |
|---|---|---|
| Treatment Only | F6SAV5 | Tubulin alpha chain |
| Treatment Only | F6ZSB4 | Tubulin Alpha-1A Chain |
| Treatment Only | F7ANC9 | Tubulin Alpha-1B Chain |
| Treatment Only | F6YRE0 | Tubulin Alpha-1C Chain |
| Treatment Only | F7D253 | UDP-Glucose 6-Dehydrogenase |
| Treatment Only | F6WHI8 | UDP-N-Acetylglucosamine Pyrophosphorylase 1 |
| Treatment Only | F6WI21 | UDP-N-Acetylhexosamine Pyrophosphorylase |
| Treatment Only | F6VDK7 | Uncharacterized Protein |
| Treatment Only | F6Y4I1 | UTP--Glucose-1-Phosphate Uridylyltransferase |
| Treatment Only | F6VSD4 | X-prolyl aminopeptidase 1 |
| Control and Treatment | F7AYK2 | 72 KDa Type IV Collagenase |
| Control and Treatment | P60708 | Actin, Cytoplasmic 1 |
| Control and Treatment | F7AAK7 | Actin, Cytoplasmic 2 |
| Control and Treatment | F6UJD4 | Adseverin |
| Control and Treatment | F7C3C6 | Aggrecan Core Protein |
| Control and Treatment | F7C450 | Alpha-2-Hs-Glycoprotein |
| Control and Treatment | F6V7C1 | Alpha-Enolase |
| Control and Treatment | Q5VI84 | Angiogenin |
| Control and Treatment | F7A971 | Angiopoietin-Related Protein 4 |
| Control and Treatment | Q8HZM6 | Annexin A1 |
| Control and Treatment | F6ZI51 | Annexin A2-Related |
| Control and Treatment | F7C0I7 | Basement Membrane-Specific Heparan Sulphate Proteoglycan Core Protein |
| Control and Treatment | Q0W9Q0 | Beta-Defensin 4A |
| Control and Treatment | F6VYL8 | Betaine--Homocysteine S-Methyltransferase |
| Control and Treatment | F6VUP8 | Betaine--Homocysteine S-Methyltransferase 1 |
| Control and Treatment | O46403 | Biglycan |
| Control and Treatment | F7C2J3 | Cartilage Intermediate Layer Protein 1 |
| Control and Treatment | F6U3D3 | Cartilage Oligomeric Matrix Protein |
| Control and Treatment | F6PQ46 | Ceruloplasmin |

| | | |
|---|---|---|
| Control and Treatment | F7AJP3 | Chitotriosidase-1 |
| Control and Treatment | F6WD70 | Chondroadherin |
| Control and Treatment | Q29482 | Clusterin |
| Control and Treatment | F6UZI2 | Coagulation Factor XIII A Chain |
| Control and Treatment | F7DE18 | Coiled-Coil Domain-Containing Protein 80 |
| Control and Treatment | F6R4Y3 | Collagen Alpha-1 |
| Control and Treatment | F6UW03 | Collagen Alpha-1 |
| Control and Treatment | F6WCC9 | Collagen Alpha-1 |
| Control and Treatment | F6Y8T1 | Collagen Alpha-1 |
| Control and Treatment | F7A3F7 | Collagen Alpha-1 |
| Control and Treatment | F6RTL6 | Collagen Alpha-2 |
| Control and Treatment | F7BRR9 | Collagen Alpha-2 |
| Control and Treatment | F7CGV8 | Collagen Alpha-2 |
| Control and Treatment | F6R735 | Collagen Alpha-3 |
| Control and Treatment | F6RVX8 | Collagen Type I Alpha 1 Chain |
| Control and Treatment | F6SSG3 | Collagen Type I Alpha 1 Chain |
| Control and Treatment | F7D939 | Collagen Type I Alpha 1 Chain |
| Control and Treatment | F7D9C7 | Collagen Type I Alpha 1 Chain |
| Control and Treatment | F6RTH9 | Collagen Type I Alpha 2 Chain |
| Control and Treatment | F6RTI8 | Collagen Type I Alpha 2 Chain |
| Control and Treatment | F6RTJ6 | Collagen Type I Alpha 2 Chain |
| Control and Treatment | F6RTN7 | Collagen Type I Alpha 2 Chain |
| Control and Treatment | F6RTP3 | Collagen Type I Alpha 2 Chain |
| Control and Treatment | F6RUA6 | Collagen Type I Alpha 2 Chain |
| Control and Treatment | F6R528 | Collagen type III alpha 1 chain |

| | | |
|---|---|---|
| Control and Treatment | F6QAT0 | Collagen type VI alpha 3 chain |
| Control and Treatment | F6TIN6 | Complement C1Q Subcomponent Subunit A |
| Control and Treatment | F7BUV8 | Complement C1Q Subcomponent Subunit B |
| Control and Treatment | F7DBT2 | Complement C1Q Subcomponent Subunit C |
| Control and Treatment | F7BTW7 | Complement C3 |
| Control and Treatment | F6XSF7 | Complement C4-A-Related |
| Control and Treatment | F6RMD0 | Complement Factor B-Related |
| Control and Treatment | F6X0K3 | Connective Tissue Growth Factor |
| Control and Treatment | F7CAX0 | C-Type Lectin Domain Family 3 Member A |
| Control and Treatment | F6WCB7 | Cytokine-Like Protein 1 |
| Control and Treatment | O46542 | Decorin |
| Control and Treatment | F7BQJ6 | Elongation Factor 1-Alpha |
| Control and Treatment | A2Q0Z0 | Elongation Factor 1-Alpha 1 |
| Control and Treatment | F6RRD0 | Elongation Factor 1-Alpha 1 |
| Control and Treatment | F6UME7 | Elongation Factor 1-Alpha 1 |
| Control and Treatment | F6UUS3 | Elongation Factor 2 |
| Control and Treatment | F6QYS3 | Extracellular Matrix Protein 1 |
| Control and Treatment | F6W2Y1 | Fibrinogen Gamma Chain |
| Control and Treatment | A2Q126 | Fibromodulin |
| Control and Treatment | F7CN05 | Fibronectin |
| Control and Treatment | F7CN11 | Fibronectin |
| Control and Treatment | F7ABC9 | Fibulin-1 |
| Control and Treatment | F6SX98 | Fructose-Bisphosphate Aldolase A |
| Control and Treatment | F6ZP69 | Fructose-Bisphosphate Aldolase C |
| Control and Treatment | Q28372 | Gelsolin |

| | | |
|---|---|---|
| Control and Treatment | F6QIB2 | Glia-Derived Nexin |
| Control and Treatment | F6YV40 | Glyceraldehyde-3-Phosphate Dehydrogenase |
| Control and Treatment | F6ZD04 | Glycogen Phosphorylase, Brain Form |
| Control and Treatment | F7AB03 | Granulin Precursor |
| Control and Treatment | F7AS83 | Granulins |
| Control and Treatment | F7DW69 | Heat Shock 70 KDa Protein 1A-Related |
| Control and Treatment | F7DMY7 | Heparan Sulfate Proteoglycan 2 |
| Control and Treatment | F6TAH8 | Hhip-Like Protein 2 |
| Control and Treatment | F6SK17 | Histone H2B Type 1-B |
| Control and Treatment | F6UGW9 | Histone H2B Type 1-J |
| Control and Treatment | F7DIN7 | Histone H2B Type 1-O |
| Control and Treatment | F7AHT9 | Histone H2B Type 2-C-Related |
| Control and Treatment | F7AZR5 | Histone H2B Type 2-E |
| Control and Treatment | F6PRL4 | Histone H2B Type 3-B |
| Control and Treatment | F7A4V1 | Histone H2B Type 3-B |
| Control and Treatment | F6VCB4 | Histone H3 |
| Control and Treatment | F6Y7V5 | Histone H3.1 |
| Control and Treatment | F7B238 | Histone H3.1T |
| Control and Treatment | F6YXV8 | Histone H3.3-Related |
| Control and Treatment | F7C964 | Histone H3-Related |
| Control and Treatment | F6VFV9 | Histone H4 |
| Control and Treatment | Q28381 | Hyaluronan And Proteoglycan Link Protein 1 |
| Control and Treatment | F7DEB1 | Insulin-Like Growth Factor-Binding Protein 6 |
| Control and Treatment | F6TNR4 | Keratin, Type I Cuticular Ha1 |
| Control and Treatment | F6SS14 | Keratin, Type I Cytoskeletal 10 |

| | | |
|---|---|---|
| Control and Treatment | F6WDW3 | Keratin, Type I Cytoskeletal 10 |
| Control and Treatment | F7ATL5 | Keratin, Type I Cytoskeletal 14 |
| Control and Treatment | F7B7X0 | Keratin, Type II Cytoskeletal 1 |
| Control and Treatment | F6W7V0 | Keratin, Type II Cytoskeletal 5 |
| Control and Treatment | F7D281 | Keratin, Type II Cytoskeletal 6B-Related |
| Control and Treatment | F6YZV8 | Keratin-87 Protein-Related |
| Control and Treatment | F7B0S3 | Lactadherin |
| Control and Treatment | F6ZDB7 | Latent-Transforming Growth Factor Beta-Binding Protein 2 |
| Control and Treatment | F6VQL3 | Leukocyte Cell-Derived Chemotaxin-2 |
| Control and Treatment | F6W3T1 | L-Lactate Dehydrogenase A Chain |
| Control and Treatment | F6SKT2 | Lumican |
| Control and Treatment | F7CU94 | Lysozyme C |
| Control and Treatment | F6S243 | Macrophage Migration Inhibitory Factor |
| Control and Treatment | F6XSR3 | Matrilin-3 |
| Control and Treatment | F6R013 | Matrix Gla Protein |
| Control and Treatment | F6T9W9 | Matrix metalloproteinase 3 |
| Control and Treatment | Q28397 | Matrix metalloproteinase 3 |
| Control and Treatment | F6VJF5 | Melanoma-Derived Growth Regulatory Protein |
| Control and Treatment | F6QKR7 | Myocilin |
| Control and Treatment | F6YY66 | Nucleoside Diphosphate Kinase B |
| Control and Treatment | A5YBL8 | Peptidyl-Prolyl Cis-Trans Isomerase B |
| Control and Treatment | F6S6J4 | Peroxiredoxin-1 |
| Control and Treatment | F6QXW2 | Phosphatidylethanolamine-Binding Protein 1 |
| Control and Treatment | F6X8Q2 | Phosphoglucomutase-1 |
| Control and Treatment | P00559 | Phosphoglycerate Kinase 1 |

| | | |
|---|---|---|
| Control and Treatment | F6ZY40 | Prelamin-A/C |
| Control and Treatment | F6WZ69 | Procollagen C-Endopeptidase Enhancer 1 |
| Control and Treatment | F6UP14 | Procollagen C-Endopeptidase Enhancer 2 |
| Control and Treatment | F6RZ46 | Prolargin |
| Control and Treatment | F6VSN9 | Protein Disulfide-Isomerase A3 |
| Control and Treatment | F6W3M5 | Pyruvate Kinase |
| Control and Treatment | F7C5F1 | Retinoic Acid Receptor Responder Protein 2 |
| Control and Treatment | F7ATC2 | Ribonuclease 4 |
| Control and Treatment | F6WNV3 | Secreted Frizzled-Related Protein 3 |
| Control and Treatment | F6T8U8 | Semaphorin-3C |
| Control and Treatment | F7B812 | Serine Protease Htra1 |
| Control and Treatment | F6ZTA7 | Serum Amyloid A Protein |
| Control and Treatment | F6ZL17 | Serum amyloid A protein |
| Control and Treatment | F7DMJ7 | Sushi Repeat-Containing Protein Srpx2 |
| Control and Treatment | F6TVT0 | Target of Nesh-Sh3 |
| Control and Treatment | F6PN81 | Tenascin |
| Control and Treatment | F7CCQ6 | Tenascin-X |
| Control and Treatment | F6YR34 | Thrombospondin-1 |
| Control and Treatment | F7E0P3 | Thrombospondin-4 |
| Control and Treatment | F6U5V3 | Transcobalamin-2 |
| Control and Treatment | F6VB94 | Transforming Growth Factor-Beta-Induced Protein Ig-H3 |
| Control and Treatment | F7D9J2 | Transketolase |
| Control and Treatment | F7DMU4 | Tumour Necrosis Factor Receptor Superfamily Member 11B |
| Control and Treatment | Q28396 | Type II collagen |
| Control and Treatment | F6SJK9 | Uncharacterized Protein |

| | | |
|---|---|---|
| Control and Treatment | F6SQQ0 | Uncharacterized Protein |
| Control and Treatment | F6TTN2 | Uncharacterized Protein |
| Control and Treatment | F6XIM5 | Uncharacterized Protein |
| Control and Treatment | F6Y2J1 | Uncharacterized Protein |
| Control and Treatment | F7CJG3 | Uncharacterized Protein |
| Control and Treatment | F7E454 | Uncharacterized Protein |
| Control and Treatment | F7B5C4 | Vimentin |
| Control and Treatment | F6QP96 | Vitrin |
| Control and Treatment | F7D8I6 | Xanthine Dehydrogenase/Oxidase |
